# Connecting structure to function with the recovery of over 1000 high-quality activated sludge metagenome-assembled genomes encoding full-length rRNA genes using long-read sequencing

**DOI:** 10.1101/2020.05.12.088096

**Authors:** Caitlin M Singleton, Francesca Petriglieri, Jannie M Kristensen, Rasmus H Kirkegaard, Thomas Y Michaelsen, Martin H Andersen, Zivile Kondrotaite, Søren M Karst, Morten S Dueholm, Per H Nielsen, Mads Albertsen

**Author notes:** Corresponding authors: Prof. Mads Albertsen & Prof. Per Halkjær Nielsen, Center for Microbial Communities, Department of Chemistry and Bioscience, Aalborg University, Fredrik Bajers Vej 7H, 9220 Aalborg, Denmark.

## Abstract

Microorganisms are critical to water recycling, pollution removal and resource recovery processes in the wastewater industry. While the structure of this complex community is increasingly understood based on 16S rRNA gene studies, this structure cannot currently be linked to functional potential due to the absence of high-quality metagenome-assembled genomes (MAGs) with full-length rRNA genes for nearly all species. Here, we sequence 23 Danish full-scale wastewater treatment plant metagenomes, producing >1 Tbp of long-read and >0.9 Tbp of short-read data. We recovered 1083 high-quality MAGs, including 57 closed circular genomes. The MAGs accounted for ~30% of the community, and meet the stringent MIMAG high-quality draft requirements including full-length rRNA genes. We show how novel high-quality MAGs in combination with >13 years of amplicon data, Raman microspectroscopy and fluorescence in situ hybridisation can be used to uncover abundant undescribed lineages belonging to important functional groups.

## Main

Since the first metagenome-assembled genomes (MAGs) were recovered in 2004^1,2^, thousands of metagenome-assembled genomes have shed light on the myriad of important functions of bacteria and archaea across the world’s ecosystems. By taking advantage of cheap short-read sequencing, compute power and new algorithms the recovery of MAGs has increased exponentially to enable over 100,000 in single studies^3^. However, MAG quality has not improved to the same extent, and there are increasing concerns regarding the quality of reference databases and the validity of research based on them^4^. The Minimum Information about a Metagenome Assembled Genome (MIMAG) standard was introduced to unify the field’s reporting standards, stating that high-quality (HQ) draft MAGs must be >90% complete, <5% contaminated and importantly include the full-length 16S, 23S, 5S rRNA genes and >18 tRNA genes^5^. Under the MIMAG standards, only a very limited number of HQ draft MAGs have been recovered from any study to date^3,6–8^. HQ reference databases are urgently required to confidently investigate the structure and function of complex microbial communities in natural or engineered ecosystems.

Wastewater treatment and resource recovery are subject to the increasing pressures of human population growth, and the demands for sustainability, human health, and reduced environmental impact. Microorganisms underpin wastewater treatment processes, from organic matter degradation and bioenergy generation to the removal of contaminants and recovery of nutrients such as nitrogen and phosphorus^9,10^. Activated sludge (AS) is the most practiced system worldwide for wastewater treatment and the functional capacity and quality of treatment is entirely dependent on the balance of the bacterial taxa within the AS biomass. Determining the structure and potential function of microbial communities in full-scale systems is central to tracking wastewater treatment efficiency and consolidating changes in structure with changes in the system, and for carrying out informed management of the communities^11^. While hundreds or thousands of MAGs have been recovered from AS systems through private and public datasets^8,12^, very few of these are HQ. HQ MAGs are beginning to be recovered from these systems^13^, and are needed to solve challenges in the treatment process. For example, an overgrowth of particular microbial morphotypes can lead to solid-liquid separation problems during the settling phase^14^, and the absence of functional groups can lead to poor nutrient recovery^15^. Here we demonstrate for the first time the combined use of long and short-read sequencing for the high-throughput production of HQ MAGs from complex microbial communities. We also present the benefits of HQ MAGs, by linking genomic functional potential to the comprehensive MiDAS3 16S rRNA gene database, with >13 years of amplicon data, used to cost-effectively monitor wastewater treatment plant (WWTP) microbial communities in Denmark^16,17^ **(Figure 1)**. This approach enables us to target, visualise and experimentally confirm the metabolic potential of an abundant yet uncharacterised genus widespread in Danish WWTPs **(Figure 1)**.

**Figure 1:**
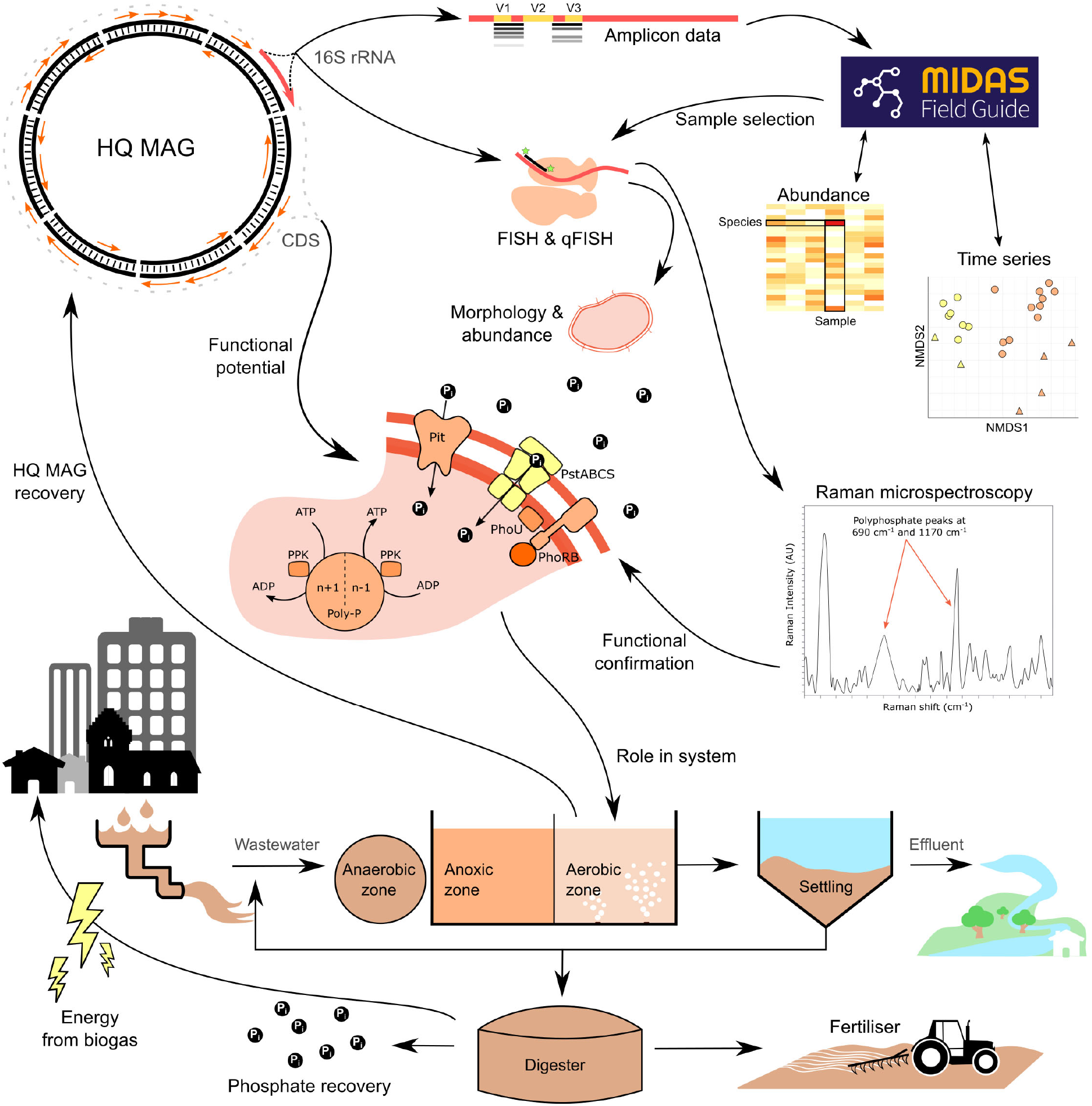
Conceptual overview of the value of HQ MAGs in linking structure to function. HQ MAGs with full-length 16S rRNA genes are recovered from a full-scale AS sample, allowing linkage to the abundance and time-series data of MiDAS and informing sample selection for further experiments. MAG and MiDAS i full-length 16S rRNA gene sequences facilitate creation of lineage specific FISH probes. Abundance is confirmed with quantitative-FISH (qFISH), and morphology and location in the floc is determined. The coding sequences (CDS) provide information on functional potential, allowing for the selection of novel species that may belong to certain functional guilds. Specific potential pathways, such as polyphosphate accumulation can be experimentally determined with Raman microspectroscopy in combination with FISH 1 and the information gained from the full-length 16S rRNA gene. This leads to confirmation of the population’s role in the AS system, and uncovers targets for investigation into improved resource recovery and effective wastewater treatment.

## Recovery of high-quality MAGs

Taking advantage of the recent gains in affordable high-throughput long-read sequencing using the Oxford Nanopore PromethlON platform, we combined 1 Tbp long-read (Oxford Nanopore) and 0.9 Tbp short-read (Illumina) data and recovered 3733 medium to HQ MAGs from across 23 Danish WWTPs **(SData1, SData2)**. Of these MAGs, 1045 meet the MIMAG standard of HQ draft genomes with >90% completeness and <5% contamination, with 19 having circularised closed metagenome assembled genomes (CMAGs) **(SData3)**. Full-length rRNA genes are usually missing in MAGs due to the difficulties associated with assembling conserved and repetitive regions using short-read sequences^18^. The combination of long and short-read sequencing methods used here enabled 91.3% (1045 out of 1145) of the near-complete MAGs to encode full-length 16S rRNA genes in addition to full-length 23S and 5S rRNA genes.

A further 38 CMAGs were identified and included in the HQ set despite not meeting the 90% completeness threshold **(SData3, Figure 2)**. These MAGs are likely complete as they were circularised and most belong to recognised streamlined or reduced genome groups, such as the bacterial lineages Patescibacteria (Candidate Phyla Radiation (CPR)) and Dependentiae^19^. However, three MAGs were from phyla not commonly associated with reduced genomes, the Proteobacteria (2 MAGs) and UBA10199 (1). A streamlined single contig genome was recently recovered for a Proteobacteria^6^, and our recovery of CMAGs for a novel Micavibrionales (2 Mbp) and Burkholderiales (0.9 Mbp) provides additional evidence of genome reduction in some members of this phylum. The group UBA10199 is undescribed and comprises a collection of only a few incomplete MAGs (24 in GTDB Release 04-RS89^20^). We believe the addition of our 38 small genome CMAGs will be valuable in revising single copy marker gene-based completeness estimates for these lineages, which are currently severely underestimated^21,22^. In total, 57 CMAGs were recovered, nearly doubling the number of CMAGs in the public domain (currently 59 reported by Chen et al. 2019).

**Figure 2:**
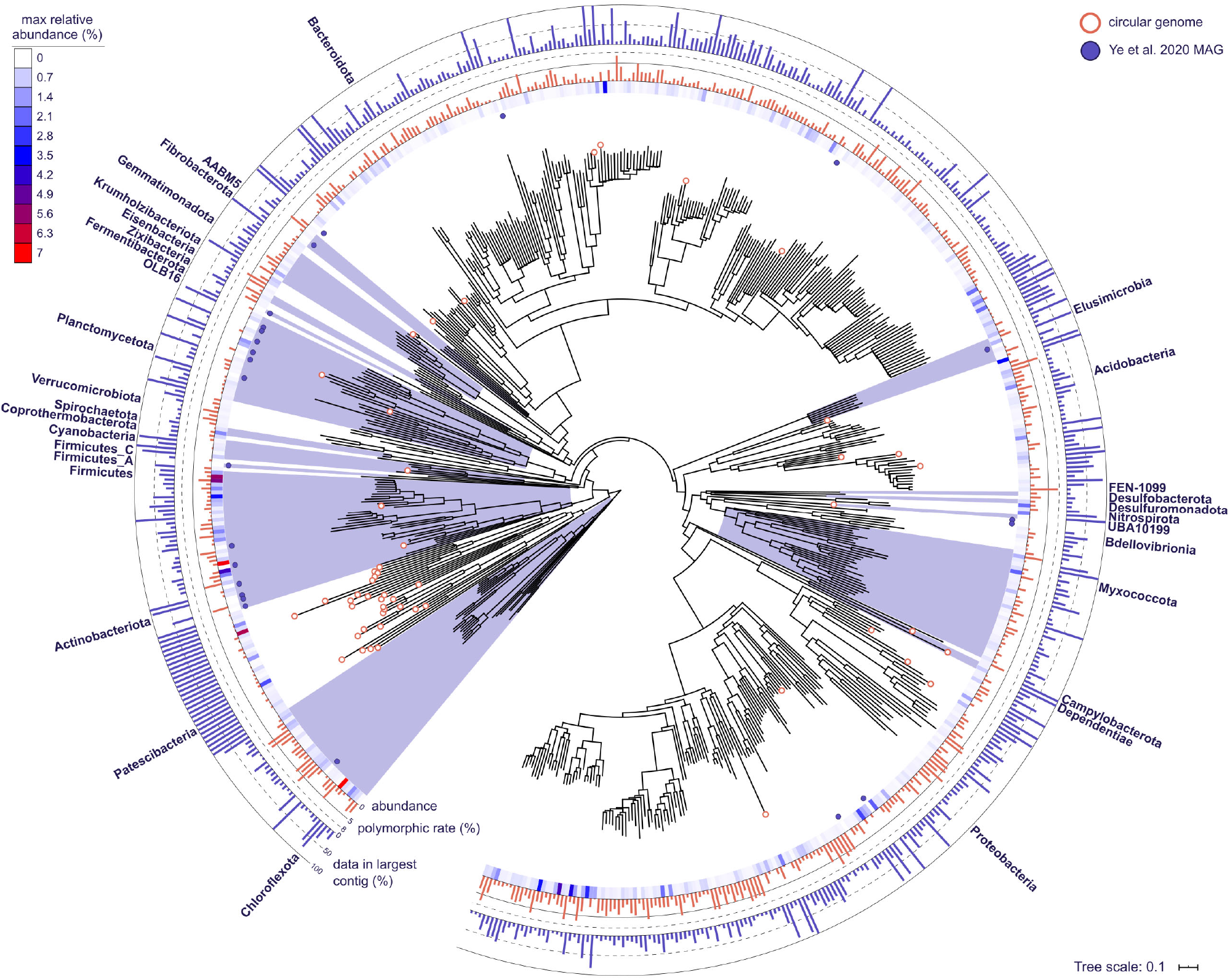
Phylogenetic bacterial genome tree based on the concatenated alignment of 120 single copy marker genes using GTDB-Tk. The 578 HQ bacterial species representatives are shown, with phyla labelled. Circular genomes are indicated at the tips using a white filled circle. HQ MAGs from Ye et al. 2020 are indicated by the purple circles. The maximum relative abundance of the MAG across the 69 WWTP metagenomes is indicated by the heatmap. Polymorphic rate is indicated by the red barchart, and percentage of data incorporated in the longest contig within the MAG is indicated in the blue barchart. Additional information on the MAGs is presented in **SData3**.

All 1083 MAGs in the HQ set encode full-length 16S rRNA, 23S rRNA, 5S rRNA, >18 tRNA genes and had polymorphic site rates ranging from 0-7.7% (average 2%, **Figure 2, SData3)**. Higher polymorphic site rates are suggestive of more chimeric population genomes and greater strain heterogeneity^3^.

## MAG diversity and community representation

The 1083 HQ MAGs represent 578 different bacterial and three different archaeal species (95% average nucleotide identity clustering) **(Figure 2, SData3)**. Phylogenetic analyses uncovered high taxonomic diversity from across 31 phyla, with the majority of MAGs attributed to the Bacteroidota (445 MAGs), Proteobacteria (276) and Chloroflexota (53). Genomes for uncharacterised phyla were also recovered, including AABM5-125-24 (4 MAGs), Krumholzibacteriota (11) and the Patescibacteria (30). An additional 317 Patescibacteria MAGs were recovered in the medium quality set, many of these are likely near-complete with 76 comprising ≤ 5 contigs. An average of 44% and 30.9% of the populations in the metagenomes were successfully assembled or binned, respectively, based on analyses of the HQ MAG single copy ribosomal protein sequences **(SFig1, SData4)**. Clustering the markers at 89% ANI showed we have recovered a HQ MAG representative for the majority (58.4%) of the genera present in the sample metagenomes **(SData4)**^7^, suggesting that we have created a representative reference database. Coverage data based on mapping metagenome reads to the 581 species supported the recovery estimates, with an average of 27.2% of sample metagenomes mapping at >95% sequence identity and >75% alignment **(SData5)**. Most MAGs had a relative abundance of ≥0.1% in at least one sample (515 species), while 66 MAGs represent consistently low relative abundance species **(SData3, SData6, Figure 2)**. Close to all of the MAGs (998 or 92%) represent populations undescribed at the species level or at higher ranks **(Figure 3a, b, SData3)**.

**Figure 3:**
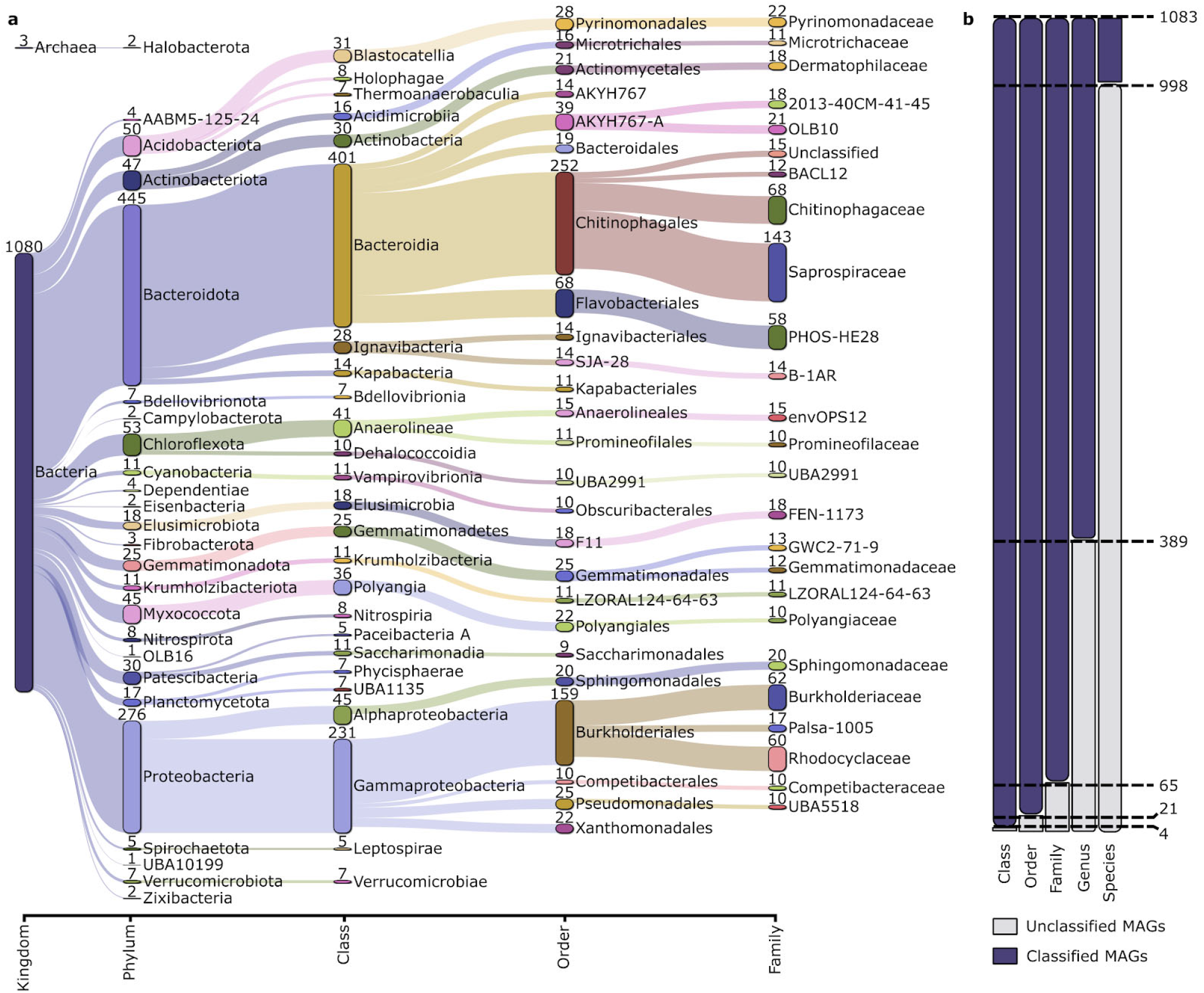
MAG recovery information across taxonomic levels, a) Sankey based on assigned taxonomy showing the novel populations at different phylogenetic levels, with the top 25 taxa shown at each level. Numbers indicate the number of MAGs recovered for the lineage, b) Total MAGs unclassified by GTDB-Tk at each taxonomic level.

Populations highly abundant in the metagenomes but not successfully binned mainly belonged to the Anaerolinea (Chloroflexota), Gamma- and Alphaproteobacteria (Proteobacteria) and Bacteroidia (Bacteroidota) **(SFig2)**. One specific Anaerolinea taxonomic unit, classified within the family envOPS12 and likely within *Ca*. Villigracilis, was consistently abundant yet unbinned across the metagenomes based on analysis of the ribosomal protein marker genes **(SFig3)**. This lineage was traced back to the MAG data, and was found in two highly contaminated and fragmented bins (192% and 250% contaminated, 86% and 95% strain heterogeneity, >500 contigs). These bins could not be manually refined due to high strain heterogeneity, which was confirmed by the presence of up to five copies of highly similar single copy ribosomal proteins in both bins **(SData7)**. Analysis of the SNPs in the single copy ribosomal protein genes confirmed the presence of multiple sub-populations of the *Ca*. Villigracilis population in each sample **(SFig4)**. This strain heterogeneity likely caused well-known downstream problems in sequence assembly and binning (Olson et al. 2019), which remains a challenge even when employing long-read sequencing^23^.

We compared our HQ MAGs to the largest AS MAG study to date by Ye et al. 2020. The authors recovered 2045 MAGs with a quality score (completeness - 5x contamination) ≥ 50 from 57 samples and 57 public datasets (114 metagenomes, including 16 metagenomes from Denmark). Of the Ye MAGs, 31% (644) met the MIMAG HQ completeness and contamination threshold (>90% completeness, <5% contamination) **(SData8)**, but only 1.2% (25 MAGs) were HQ according to the MIMAG standards and encoded the rRNA genes **(SData9, Figure 2)**. After 95% ANI clustering with the HQ species representatives of this study, the 25 HQ Ye MAGs were found to belong to 24 species from predominantly the Planctomycetota (6 MAGs) and Actinobacteriota (6 MAGs) phyla **(SData10)**. Three species, *Microthrixparvicella* and two MAGs belonging to uncharacterised lineages in the Elusimicrobia and Actinobacteriota, were represented by a better quality MAG from our set **(SData10)**. The benefits of using long-read sequencing to improve both MAG quality and quantity are clear as we recovered over 40x more HQ MAGs from fewer (23 vs 114) metagenomes and a similar amount of data (830 Gb long-read 922 Gb short-read, vs 1352 Gb). Using this method for large scale recovery efforts in similarly complex environments, such as soil and the human gut, would lead to huge increases in the quality of all associated MAG reference databases.

However, it is important to acknowledge the widespread limitations of MAGs that represent chimeric population bins (not strains), are recovered in pipelines without individual detailed manual curation, and are affected by sequencing platform errors and low coverage regions^13,21^. While still imperfect, this comprehensive collection of HQ MAGs provides the first links between 16S rRNA gene based studies and the complete gene repertoires for many uncharacterised organisms in AS.

## Functional guilds central to wastewater resource recovery

The recovered MAGs facilitated analysis into the functional potential of the microbes involved in important wastewater processes such as nitrification, denitrification, enhanced biological phosphate removal (EBPR), floc formation and solid-liquid separation. Removal of ammonium and other nitrogen species from wastewater is essential to prevent pollution and ecosystem disturbances^24,25^. Both nitrifiers and denitrifiers are required for efficient removal, by oxidising ammonia and/or nitrite to nitrate, and then reducing nitrate to nitrogen gas. Ammonia oxidisers were represented by seven *Nitrosomonas* MAGs belonging to six different species with only one, *N. oligotropha*, characterised **(SData3)**. In addition, three *Nitrospira* comammox MAGs were obtained, including one novel species and two *N. nitrosa*. The nitrifiers also included five MAGs belonging to the same species as *Nitrospira* sp. strain ND1, related to *N. defluvii* (~92% ANI)^26^, and two *Nitrotoga* MAGs belonging to a new species (94% ANI to *Candidatus* Nitrotoga sp. CP45^27^). These results show that a large undescribed diversity remains hidden even within well-studied functional guilds such as nitrifiers, for which genomes will be valuable in future targeted studies.

Nitrate appeared to be widely used as a potential electron acceptor as many of the MAGs (214) encoded the nitrate reductase (NarGHI) **(Figure 4, SFig5)**. Denitrifiers encode the complete pathway for the reduction of nitrate to nitrogen gas, and 21 MAGs were capable of denitrification based on the genome annotations **(SData11)**. Well known denitrifiers in AS are often found within the Gammaproteobacteria, such as *Azoarcus, Dechloromonas, Thauera* and *Zoogloea*^28^. Similarly, most of the identified denitrifiers in the HQ MAG set belonged to the Gammaproteobacteria (15 MAGs), and included three additional genomes of *Zoogloea* belonging to two novel species and eight *Dechloromonas*. Interestingly, nearly half of the MAGs (489) encoded NosZ, which catalyses the reduction of N_2_O to N_2_ **(Figure 4)**, suggesting that non-denitrifiers are important in N_2_O reduction and potentially mitigate the release of this greenhouse gas to the environment 29

**Figure 4:**
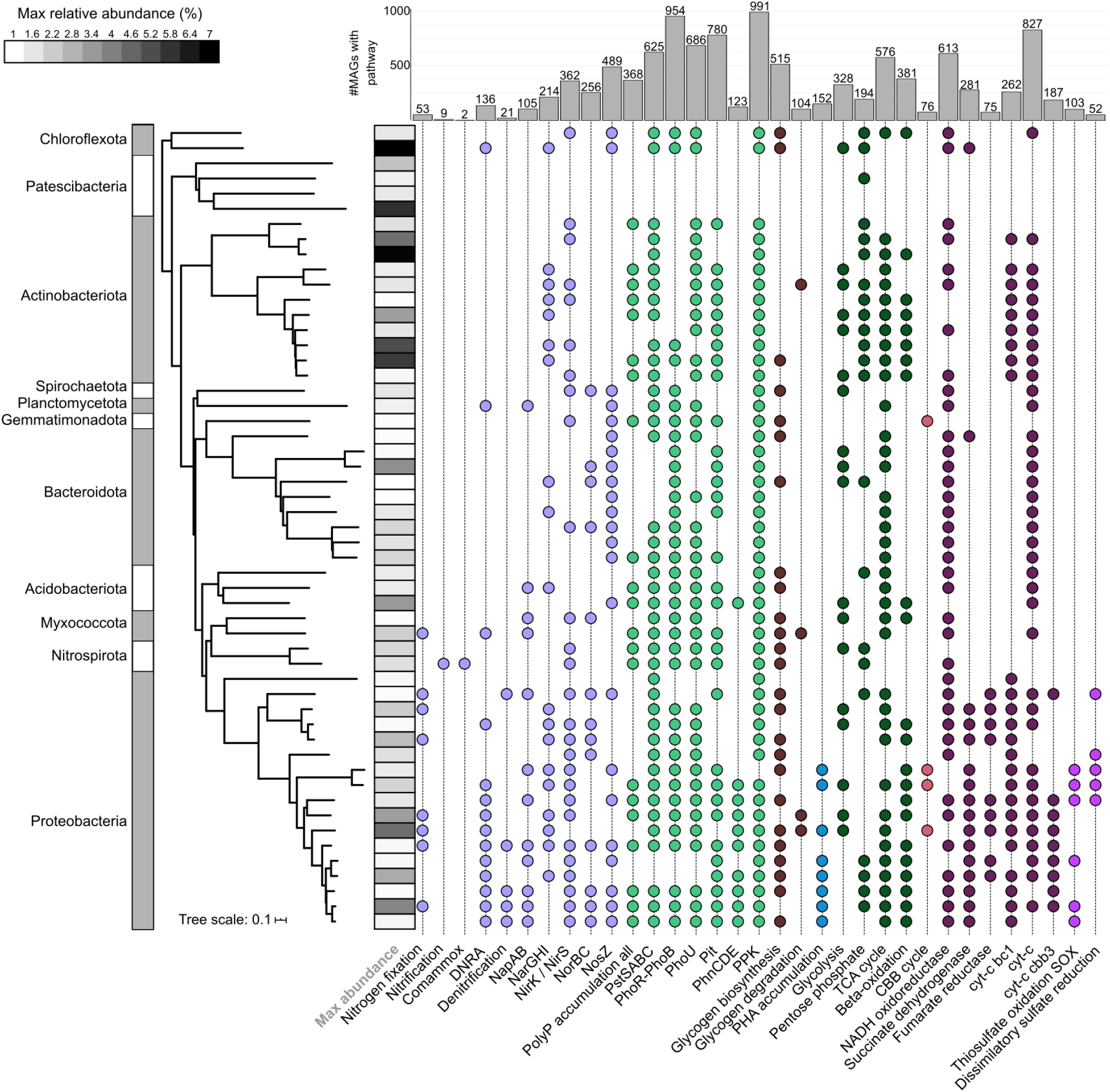
Functional profiles of the top 53 bacterial species representatives with relative abundances >1% in at least one sample metagenome used in this study. Pathways are considered present if 100% of the genes in the KEGG module, or custom module **(SData12)**, are encoded. Heatmap strip indicates the maximum relative abundance of the population in the metagenomes. Bar chart indicates the number of MAGs encoding the pathway of interest. **SFig5** shows the full taxonomic string for the nodes.

In contrast to denitrification, dissimilatory nitrate reduction to ammonium (DNRA) is undesired in wastewater treatment as nitrate is converted back to ammonium rather than being removed from the system as nitrogen gas^30^. Further, DNRA microorganisms compete with denitrifiers for nitrate, and can also compete for similar electron donors. Potential DNRA (NarGH/NapAB, NrfAH) microorganisms using nitrite in respiration were identified in 43 MAGs **(SData11)**. *Geothrix* spp. enriched from AS have been found to perform DNRA when fed acetate^31^, and three *Geothrix* MAGs (2 spp.) were recovered with the genomic capacity. Two *Anaeromyxobacter* MAGs also encoded the capability, which has been identified in the soil *bacterium Anaeromyxobacter dehalogenans*^32^. Additional lineages encoding DNRA genomic capabilities included two *Rubrivivax* spp. (7 MAGs, *Leptothrix* MiDAS3 tax), and a novel *Ca*. Amarolinea species (4 MAGs). Potential use of the non-respiratory NirBD in DNRA was identified in numerous *Ca*. Accumulibacter, *Dechloromonas, Propionivibrio* and *Ca*. Competibacter MAGs **(SData11)**.

The metagenomes were derived from WWTPs performing EBPR in addition to nitrogen removal, consequently the polyphosphate-accumulating organisms (PAOs) that are responsible for this process are of key interest. Of the well-known and confirmed PAOs, we recovered *Tetrasphaera* (14 MAGs MiDAS3 tax), and *Ca*. Accumulibacter (5 MAGs). Phosphate transporter and regulation genes (*pstABCS,pit* and*phoU*) were identified in 368 MAGs in total **(SData11, SData12, Figure 4)**. The presence of these genes does not unequivocally identify PAOs, as many microorganisms can accumulate polyphosphate without cycling it^33–35^. However, these indications of potential can be used to guide *in situ* tests or enrichments in order to identify novel true PAOs (see below).

Glycogen accumulating organisms (GAOs) are historically believed to compete with PAOs for substrates and have similar genome characteristics, and in some cases similar phylogenies, to PAOs. However, GAOs do not cycle phosphate and instead use glycogen as a storage compound for use in anaerobic conditions^33^. Recent research indicates that GAOs are likely not detrimental to the activity of PAOs in full-scale systems but are abundant members of the community with many other functions^36^. Many genomes were recovered for known GAOs such as *Defluviicoccus* (3 MAGs MiDAS3 tax), *Micropruina* (3 MAGs), *Ca*.

Competibacter (7 MAGs), *Contendobacter* (3 MAGs) and *Propionivibrio* (10 MAGs)^37^, and will be valuable for confirming genus level variation in functionality. The complete pathway for glycogen biosynthesis and degradation was encoded in 79 MAGs in total **(SData11, Figure 4)**, reinforcing knowledge that this storage polymer is common among AS microorganisms^37^.

Many PAOs and GAOs rely on the carbon storage compound polyhydroxyalkanoates (PHA) when conditions become anaerobic during EBPR^35^. We identified the required metabolic pathway in 152 MAGs **(Figure 4)**, including the experimentally validated PHA-accumulating AS populations *Dechloromonas* (8 MAGs), *Ca*. Accumulibacter (5 MAGs), *Propionivibrio* (10 MAGs), and *Zoogloea* (3 MAGs)^38,39^. Excluding *Austwickia* (Actinobacteriota), additional unknown but potential PHA-accumulating populations were all from the Alpha- and Gammaproteobacteria, and predominantly belonged to the genera *Rubrivivax* (13 MAGs), *Rhodoferax* (15 MAGs) and UBA 1936 (10 MAGs).

Microbial morphology, such as filamentous growth, has a large impact on wastewater treatment efficiency as specific populations are essential for flocculation, settling and dewatering in AS, but overgrowth can lead to problems in sludge settleability (bulking) and foaming^40^. This leads to solids carryover and reduction in effluent quality, with downstream negative effects on both environmental and human ecosystem health^40^. Numerous filamentous taxa were recovered in the HQ MAG set, including the key groups *Ca*. Microthrix (10 MAGs), *Ca*. Promineofilum (1 MAG), *Ca*. Villigracilis (14 MAGs MiDAS3 tax), *Ca*. Amarolinea (4 MAGs MiDAS3 tax) and *Leptothrix* (9 MAGs MiDAS3 tax) **(SData3)**.

The recovery of 347 MQ Patescibacteria MAGs of which 30 were circular suggested a prevalence and likely functional importance of this group not previously recognised in AS. Saccharimonadia lineages previously described and visualised using FISH were consistently abundant^41,42^, but additional Patescibacteria groups were also found in high abundances of up to 5.5% **(Figure 1)**. This included two MAGs from the class Paceibacteria, belonging to different families, with 5% (closed CMAG) and 2% (single contig) relative abundances in the WWTP metagenome of Esbjerg **W (SNotel)**. The highly abundant Paceibacteria were visualised with FISH, using probes developed from the recovered MAG full-length 16S rRNA genes **(SFig6, STable1)**. Cells were found to be ultrasmall (<0.2 μm), similar to many Patescibacteria lineages^19^, and were visualised in two samples. In Esbjerg W, where an abundance of >7% was determined in the metagenome suggesting a bloom event **(SData3)**, they appeared as widespread consistent clusters, whereas in Viborg (abundance 0.7%), they appeared as isolated clusters edging the AS flocs **(SFig7)**. The Patescibacteria are believed to be primarily host-dependent as either syntrophs or parasitic populations^43^, and based on our FISH analyses this Paceibacteria lineage could potentially target multiple hosts. Overall, the abundances and diversity of the Patescibacteria suggest they likely have significant involvement in AS microbial dynamics warranting further investigation.

## Linking HQ MAGs to amplicon data

Application of the MIMAG HQ benchmark would greatly benefit the validity of investigations into uncultured microorganisms through reduced chimerisms in bins, better gene syntenic information due to improved assembly contiguity, and the recovery of multicopy genes and conserved single copy genes that are normally missing in short-read assemblies^22^. In our case, using HQ MAGs with full-length 16S rRNA genes allowed us to connect the recovered MAGs to amplicon sequencing data from the MiDAS3 project^17^, thus fulfilling the essential purpose of the database: to link function to long-term structure trends and process data **(Figure 1)**. The 13 years of amplicon data from 20 WWTPs allowed for the identification of the consistently abundant and widespread populations, and prospective target microorganisms for closer investigation. The MAG 16S rRNA genes were mapped to the MiDAS3 full-length rRNA gene database and taxonomy, which revealed we had recovered HQ representatives for 65 of the top 100 species found in WWTP across Denmark **(SData13)**^17^. At the ASV level, providing the highest level of resolution for amplicon data and a potential for species-level resolution 58 of the top 100 most abundant ASVs (averaged across MiDAS3) were recovered in a HQ MAG **(SData14)**. These recovered populations are not only abundant in Danish systems, but likely represent important populations in nutrient removal plants across the world^17,44^.

The 16S rRNA gene amplicon data was linked to the MAG metabolic annotations in order to investigate novel and consistently abundant populations with metabolisms relevant to resource recovery and bioremediation, such as the polyphosphate, PHA, glycogen and denitrification metabolisms described above. Based on the MAG 16S rRNA gene sequences and previous efforts to recover full-length 16S rRNA gene sequences from AS^17,44^, FISH probes were designed to target a potential polyphosphate-accumulating population belonging to a novel genus, midas_g 190, most closely related to *Methyloversatilis* (<78% ANI) **(STable1, SNote2)**. Phylogenetic analysis of these sequences revealed an evident separation into three species, providing for the first time an overview of the novel genus **(SFig8)**. Application of the newly designed probes revealed 0.8-1 x 0.5-0.6 μm rod-shaped cells that were often arranged in microcolonies inside the flocs, or sometimes attached to filaments **(Figure 5a)**. The relative abundances determined by amplicon sequencing were in the same range as quantitative-FISH **(STable2)**. Two species-level MAGs were recovered from within this genus (4 MAGs midas_s_484, 1 MAG midas_s_514). Using Raman-FISH microspectroscopy for the analysis of intracellular storage polymers, the genus was experimentally determined to store polyphosphate at levels similar to known PAOs, confirming the metabolism observed in the genomes **(Figure 5b,c, SData15)**, and suggesting this population may have an important role in phosphate recovery in the AS system. No Raman peaks were found for the other intracellular storage compounds glycogen and PHA. Due to genomic and experimental evidence for phosphate accumulation, and the genomic potential of methylotrophy combined with relatedness to a characterised methylotrophic genus, we propose the names *Ca*. Methylophosphatis haderslevensis (midas_s_514) and roskildensis (midas_s_484) for the two species-level populations.

**Figure 5.**
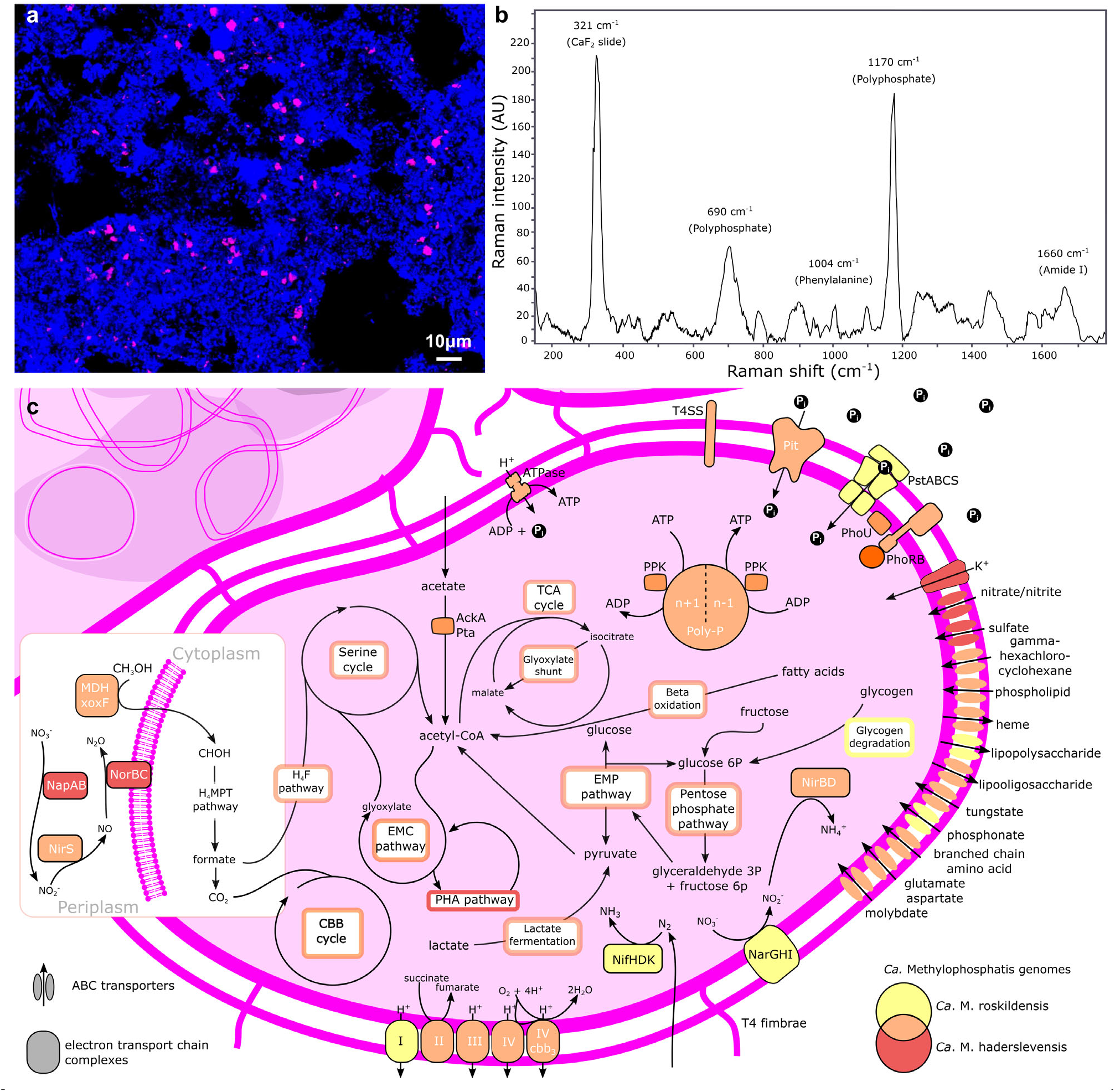
a) FISH micrograph of *Ca*. Methylophosphatis, targeted by the genus-specific probe g190_1276 (Cy3-labeled) in full-scale activated sludge from Bjergmarken WWTP. Target cells appear magenta, while all other bacterial cells appear blue, b) Raman spectrum of *Ca*. Methylophosphatis (average of 100 FISH-defmed cells) showing the presence of the signature peaks for polyphosphate (690 and 1170 cm^-1^). Peaks for phenylalanine (1004 cm^-1^) and amide I linkages of proteins (1450 cm^-1^) are specific markers for biological material, c) Metabolic reconstruction of the *Ca*. Methylophosphatis MAGs. Colours represent the species or combination of species (Venn diagram) that encode the potential for the enzyme or cycle. Abbreviations: H_4_MPT = tetrahydromethanopterin pathway, H_4_F = tetrahydrofolate pathway, TCA = tricarboxylic acid cycle, EMC = ethylmalonyl-CoA pathway, EMP = Embden-Meyerhof-Pamas pathway (glycolysis), CBB = Calvin-Benson-Bassham cycle, PHA = polyhydroxyalkanoate pathway, nitrogenase (NifHDK), CH_3_OH = methanol, methanol dehydrogenase (MDH-xoxF), I = complex I NADH dehydrogenase, II = complex II succinate dehydrogenase, III = complex III cytochrome bc1, ΓV = cytochrome c oxidase, IV cbb3 = complex IV cytochrome cbb3 oxidase, inorganic phosphate transporter family (Pit), inorganic phosphate ABC transporter (PstABCS), two component system for phosphate regulation (PhoRB), phosphate transport system accessory protein (PhoU), Poly-P = polyphosphate, type IV secretion system (T4SS), type IV fimbriae (T4 fimbriae), nitrate reductase respiratory (NarGHI), periplasmic nitrate reductase (NapAB), nitrite reductase (NirS), nitric oxide reductase (NorBC), acetate kinase (AckA), phosphotransacetylase (Pta).

## Conclusion

Here we present a methodology for the high throughput production of HQ MAGs and its application to investigate complex microbial communities, focusing on the AS system. This approach enabled amplicon and full-length 16S rRNA gene data to be linked to the functional potential of entire HQ MAGs. We show that producing HQ MAGs from complex environments is increasingly an affordable and feasible undertaking. Raising MAG standards facilitates the identification and experimental confirmation of key functional microbes, demonstrated by our characterisation of the polyphosphate accumulating novel genus *Ca*. Methylophosphatis. Furthermore, raising standards will also aid exploratory studies by preventing contamination of public repositories with low quality MAGs^4^. We expect that the rate of improvement in DNA extraction methods as well as long-read sequencing technologies indicates that thousands of HQ MAGs and CMAGs are on the horizon.

## Supporting information

SData1

SData2

SData3

SData4

SData5

SData6

SData7

SData8

SData9

SData10

SData11

SData12

SData13

SData14

SData15

## Supplementary Figures

**SFig1:**
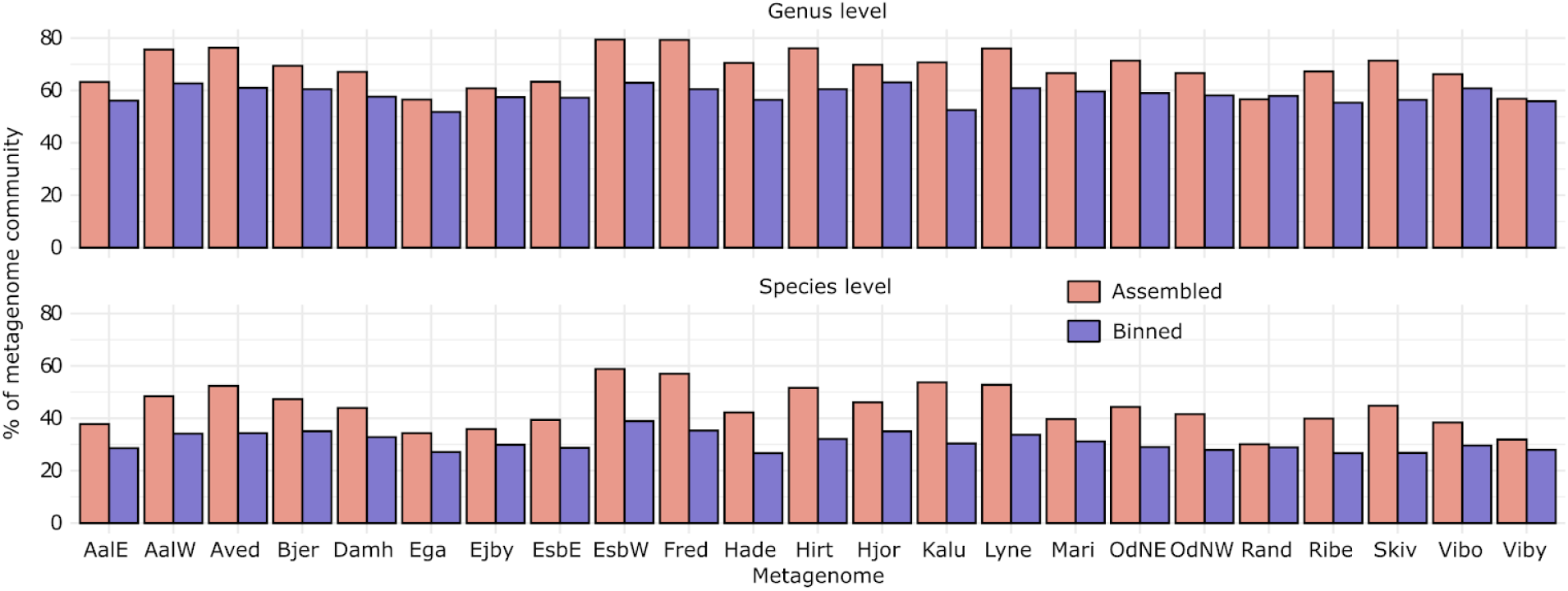
Results of the single copy ribosomal protein analysis from SingleM, showing the proportion of the metagenome populations that were successfully assembled and binned per WWTR Abundance is determined from the single copy ribosomal protein gene sequences, as taxonomic units.

**SFig2:**
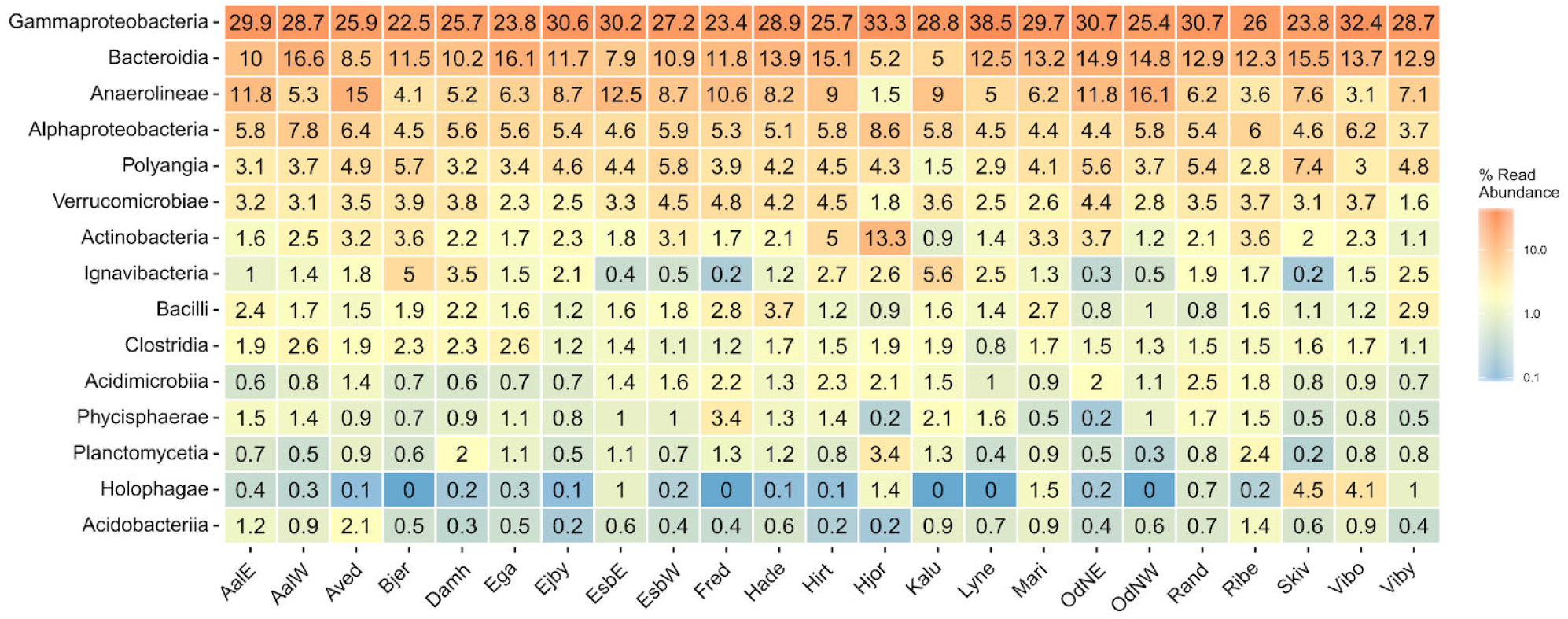
SingleM unbinned OTUs distribution heatmap showing the top 15 classes for the marker gene *rplE* for ribosomal protein L5. Abundance is relative to the total unbinned marker gene sequences.

**SFig3:**
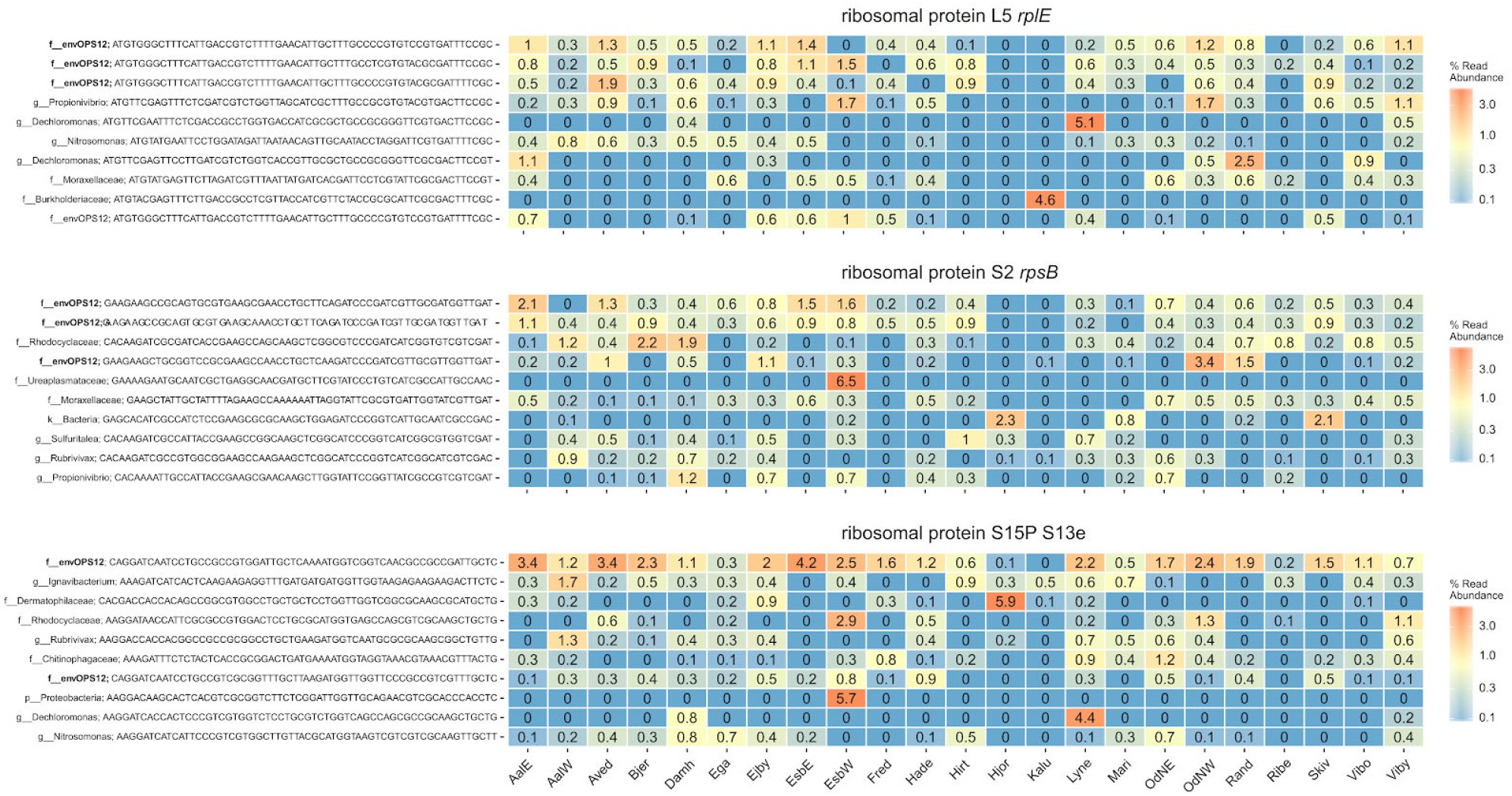
SingleM unbinned taxonomic units (representing discrete populations) distribution heatmap of the top 10 populations for three ribosomal protein marker genes. The envOPS12 populations are bolded. Abundance is relative to the total unbinned marker gene sequences.

**SFig4:**
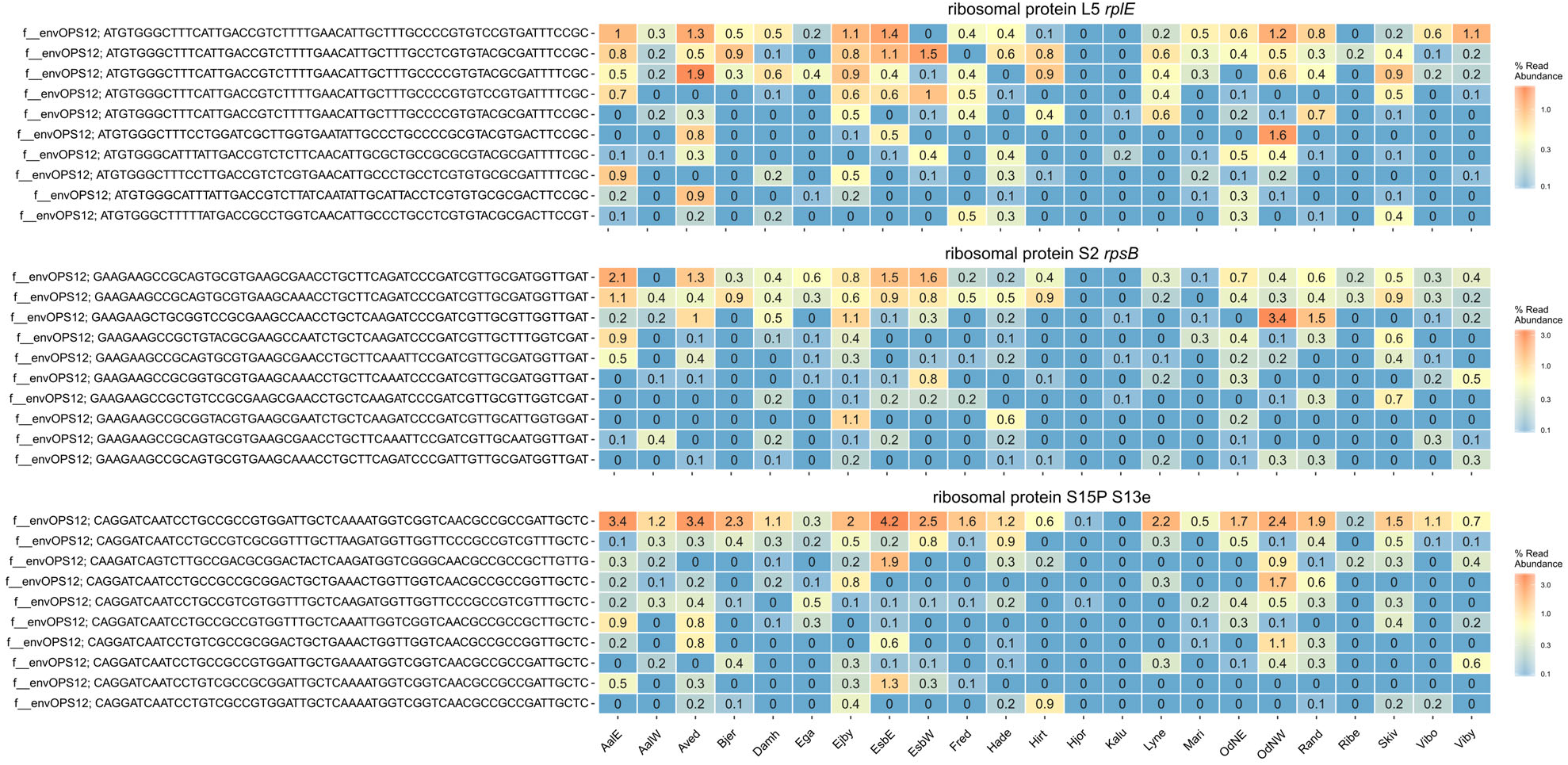
SingleM unbinned envOPS12 taxonomic unit distribution heatmap of the top 10 populations for three ribosomal protein marker genes. Abundance is relative to the total unbinned marker gene sequences.

**SFig5:**
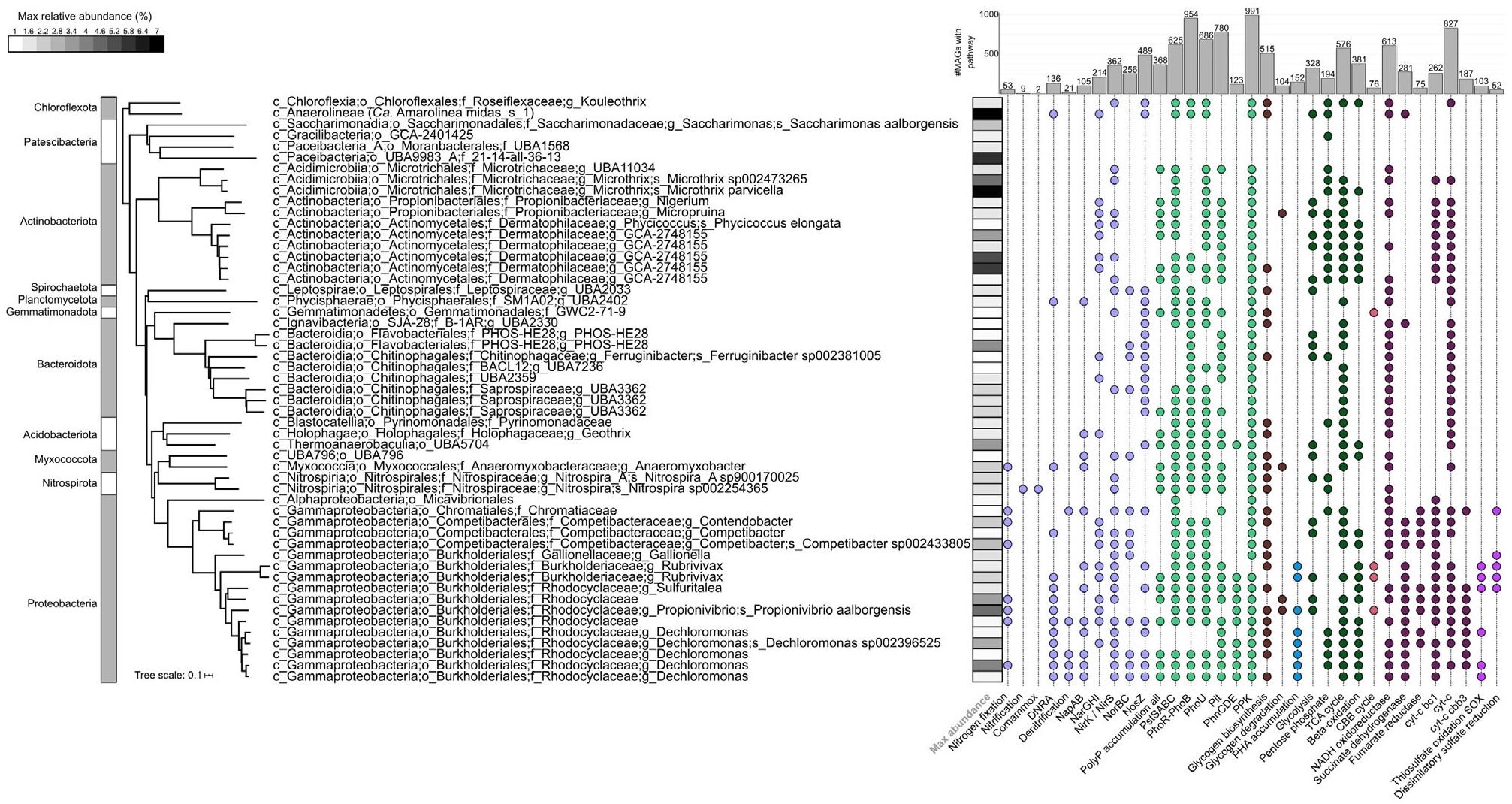
Functional profiles of the top 53 bacterial species representatives with relative abundances >1% in at least one sample metagenome used in this study. Taxonomy strings from GTDB are shown. Pathways are considered present if 100% of the genes in the KEGG module, or custom module **(SData12)**, are encoded. Heatmap strip indicates the maximum relative abundance of the population in the metagenomes. Bar chart indicates the number of MAGs encoding the pathway of interest.

**SFig6:**
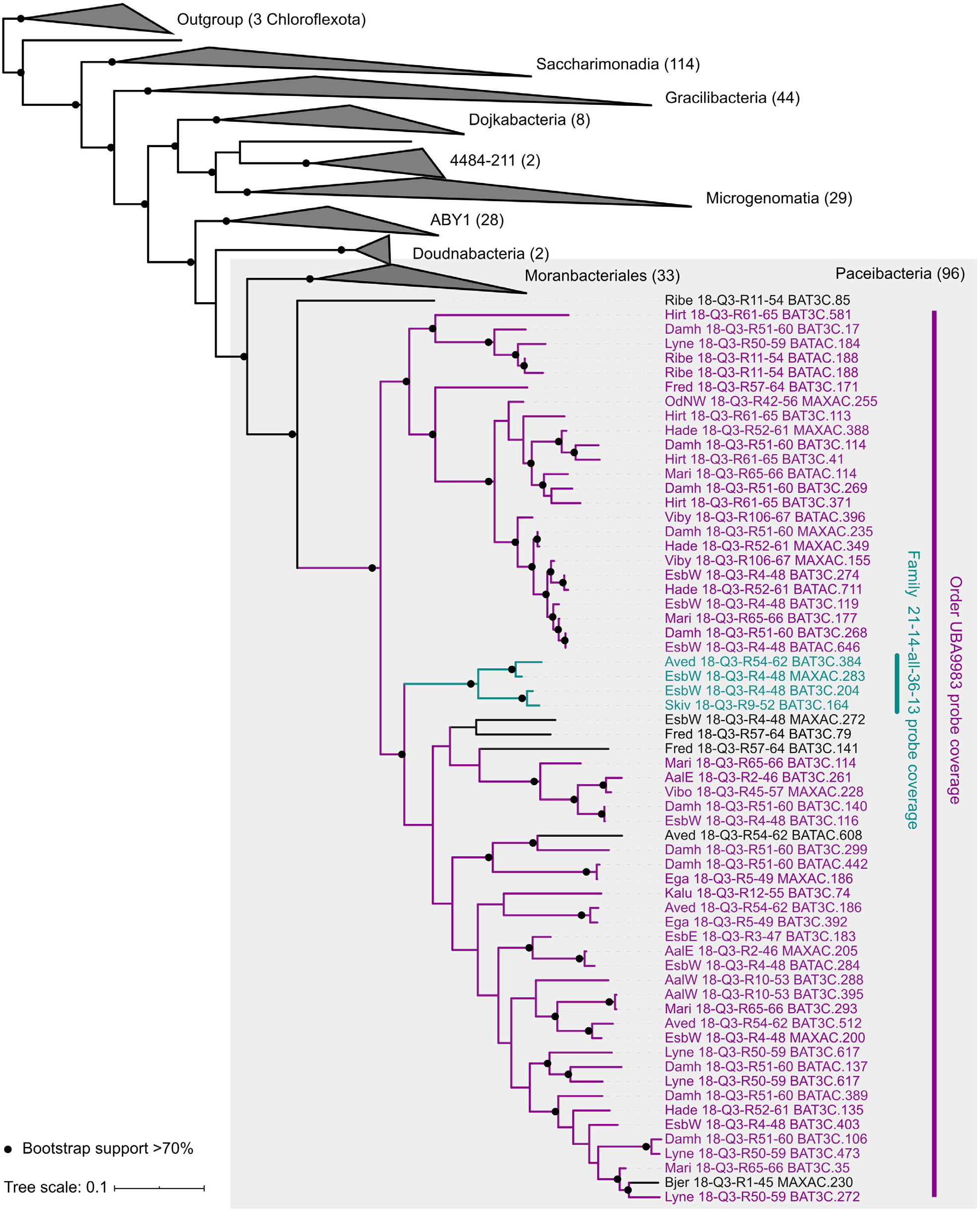
Maximum likelihood phylogenetic 16S rRNA gene tree of Patescibacteria MAG sequences, indicating the coverage of the two probes used in the FISH analyses. MAGs in black are missed by the probe set. The family probe (magenta) and order probe (cyan) overlap. Tree was bootstrapped 100x.

**SFig7:**
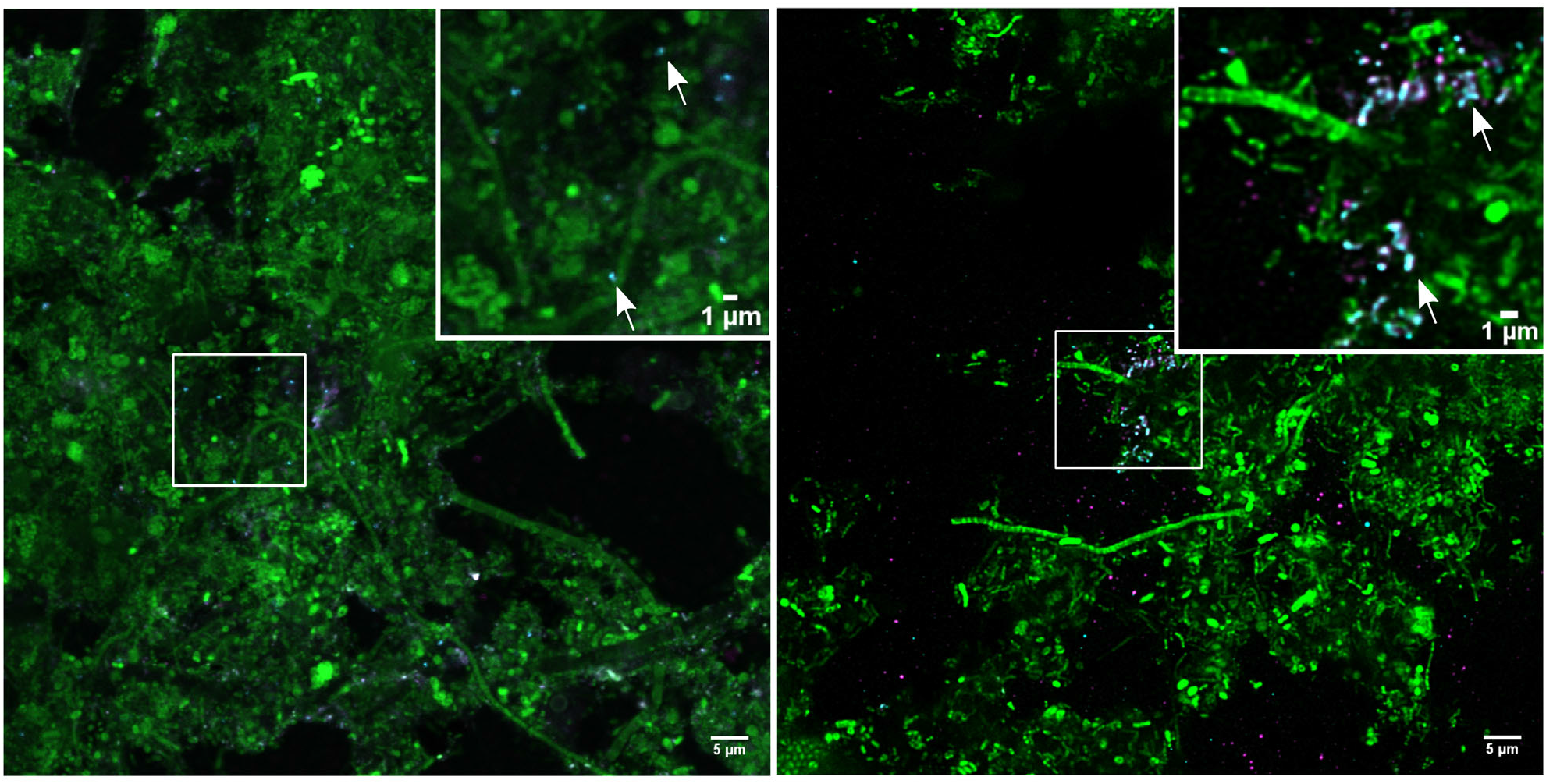
FISH images showing the ultra-small (<0.2 μm) Paceibacteria lineage. Cyan indicates the family level' probe (ATTO 532) and magenta the ‘order level’ probe (ATTO 594), and green indicates the other bacteria (EUB mix ATTO 633). Paceibacteria were very abundant (>7% based on metagenome relative abundance) and found as dispersed cells in the Esbjerg W AS sample (left), and localised cell clusters in lower abundance (0.7%) in the Viborg AS sample (right). Insets show magnified areas, arrows indicate cells.

**SFig8:**
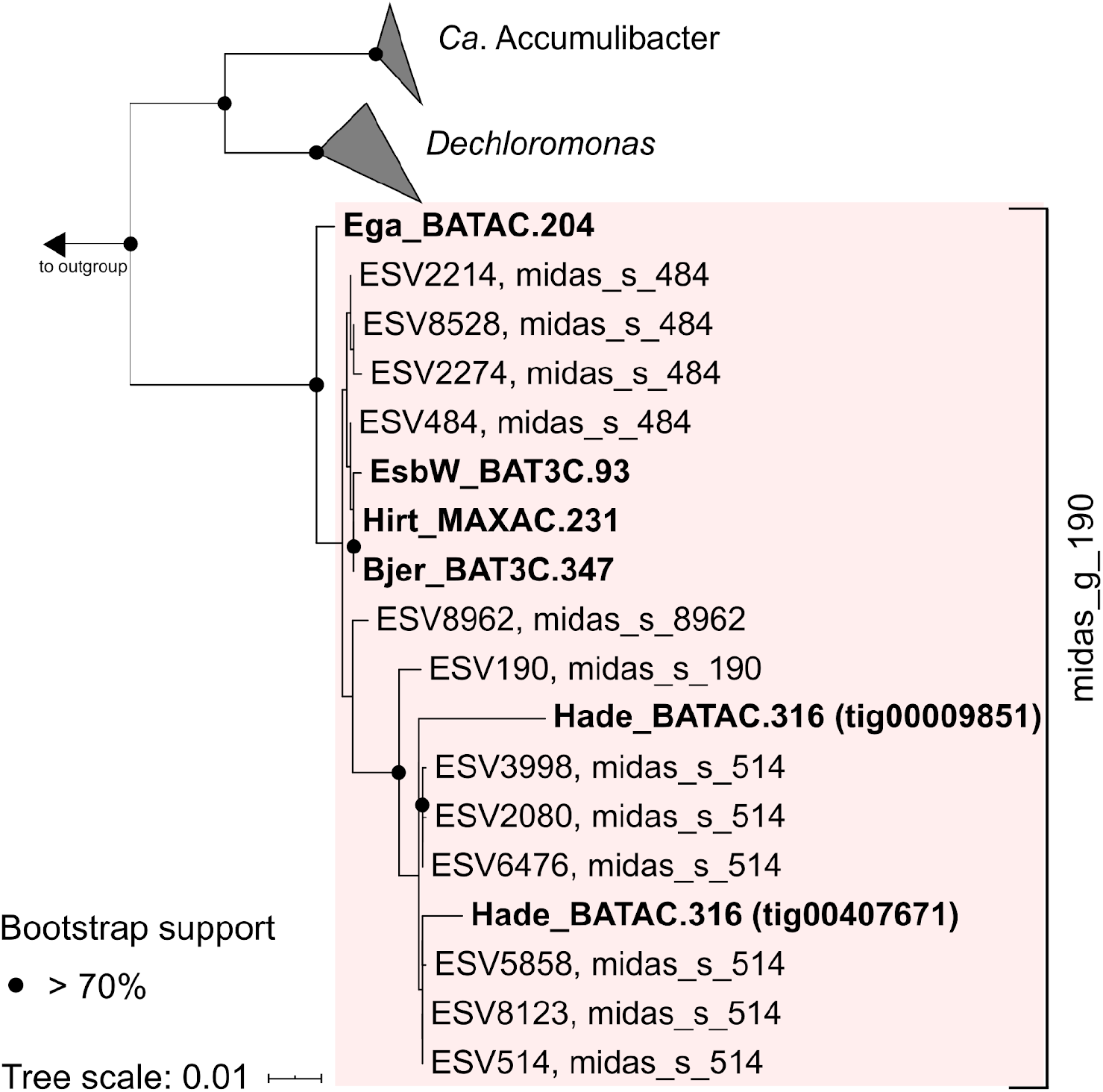
Maximum likelihood tree of the 16S rRNA gene sequences from the midas_g_190 MAGs and MiDAS3 reference database. 1000-replicate bootstraps were used. The orange box is indicating coverage of the genus-specific probe g190_1276.

## Supplementary Data

**SData1:** Data yield for each metagenome and associated sample metadata

**SData2:** Flow cell yield and summary statistics for the PromethION run

**SData3:** MAG summary statistics including: full-length rRNA genes, MiDAS3 taxonomy, SILVA v138 taxonomy and GTDB taxonomy, number of contigs, total base pairs, species representatives, completeness, contamination, polymorphic rate, primer mismatch, and coverage in the illumina and nanopore data of origin.

**SData4:** Single copy markers in MAGs and assemblies identified using SingleM, for the 23 samples used for the individual sample assemblies.

**SData5:** Coverage based abundances (CoverM) of MAGs across all 69 metagenomes.

**SData6:** Relative abundances of the 581 species representatives across the 69 Illumina metagenomes.

**SData7:** Redundant single copy ribosomal protein marker genes in envOPS12.

**SData8:** Ye et al. 2020 MAG completeness and contamination results from CheckM.

**SData9:** Identified Ye et al. 2020 MAG full length rRNA genes.

**SData10:** Ye et al. 2020 MAG dRep species dereplication with the MAGs from this study.

**SData11:** 100% complete KEGG module paths and associated KOs from EnrichM.

**SData12:** Custom modules used for EnrichM analysis, and pathways used in Figure 4.

**SData13:** Top 100 MiDAS3 species to MAGs.

**SData14:** Top 100 MiDAS3 ASVs to MAGs.

**SData15:** Specific KO hits for *Ca*. Methylophosphatis metabolic reconstruction.

## Supplementary Tables

**STable1:**
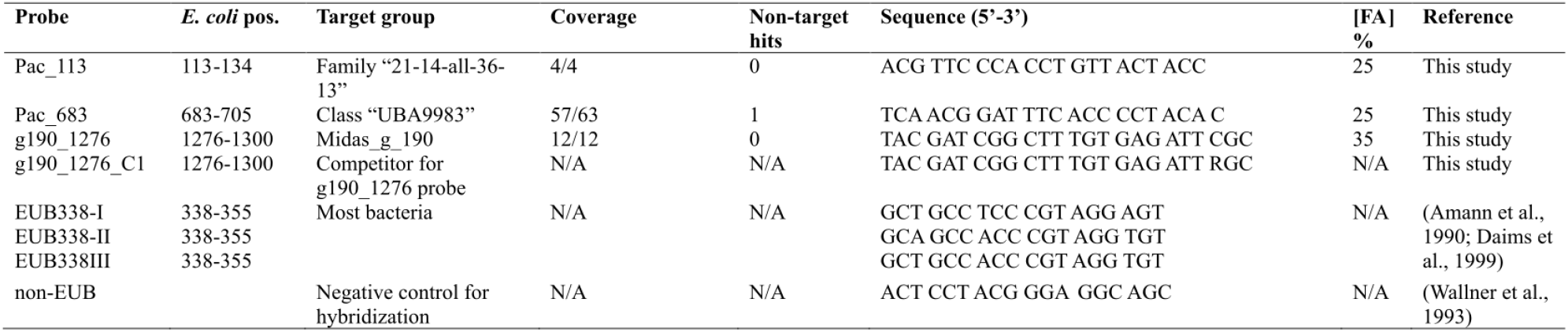
Summary of FISH gene probes used for Paceibacteria and *Ca*. Methylophosphatis (midas_g_190). Coverage for Paceibacteria 16S rRNA sequences is from the genomes only, as the target group is not available in MiDAS3. Only 1 non-target hit for Pac_683 was determined in MiDAS3. Pac_683 targets 508 sequences in SILVA v138.444 of these sequences are from within the Parcubacteria, the remaining non-target bacteria hits are within the Patescibacteria (Candidate Phyla Radiation). Pac_133 has 0 non-target hits in SILVA v138, and targets one sequence from the Cambellbacteria. Pac_113 was used with ATTO 532, Pac_683 in ATTO 594 and EUBmix in ATTO 633 to ensure no crosstalk between the dyes.

**STable2:** Summary of amplicon sequencing and qFISH relative abundances of *Ca*. Methylophosphatis (midas_g_190).

| WWTP | Sample date | Abundance (%) |  |
| --- | --- | --- | --- |
|  |  | Sequencing | qFISH |
| **Midas_g_190** |  |  |  |
| Bjergmarken | August 2018 | 1 | 2.2 ± 0.7 |
| Hjørring | August 2015 | 1 | 1.2 ± 0.6 |
| Ribe | August 2013 | 2.3 | 2.3 ± 0.9 |
| Ringkøbing | June 2014 | 1.6 | 2.2 ± 0.7 |

## Supplementary Notes

### Supplementary Note 1

Initially the most abundant Paceibacteria population (MAG EsbW_18-Q3-R4-48_MAXAC.283) was recovered in two contigs. Extraction and mapping of the Nanopore and Illumina reads belonging to the MAG, followed by reassembly with Flye enabled the MAG to be closed. Examination of the mapping data and assembly graph showed that the MAG could be circularised either with, or without, a 40 kbp DNA fragment. This fragment had 50% less coverage than the rest of the genome, and was found to insert between a tandem repeat in the population genome. The 40 kbp fragment encoded 43 CDS regions, predominantly hypothetical proteins as well as phage cluster methyltransferases, DNA methylases, phage portal, peptidase and capsid proteins. Both VirSorter v1.0.5 (category 2 prophage) and PHASTER identified the insert as viral^45,46^. Only three bp were identified as potentially polymorphic **(SData3)**, indicating the presence of a single dominant population and little strain heterogeneity. Consequently, the use of long-read sequencing enabled us to determine that half of this Paceibacteria population appears to be infected by an active phage system.

### Supplementary Note 2

The *Ca*. Methylophosphatis populations encode a versatile metabolism, including the methylotrophy pathway. Based on the location of this new genus next to the well characterised methylotrophic genus *Methyloversatilis*, we suggest methylotrophy is a potential strategy for energy generation and/or carbon assimilation. C1 carbon dissimilation was indicated by presence of the XoxF methanol dehydrogenase in four of the five MAGs (both species), as well as the tetrahydromethanopterin pathway for formaldehyde oxidation to formate, and formate oxidation genes **(SData15)**. C1 assimilation to acetyl-CoA was encoded by the tetrahydrofolate pathway and key genes of the serine cycle **(SData15)**. Furthermore, carbon assimilation could also occur through the Calvin-Benson-Bassham cycle **(Figure 5, SData15)**. However, unlike *Methyloversatilis* spp., the MAGs encoded no pathway for methylamine use (methylamine dehydrogenase MauA, MauB, or methylglutamate dehydrogenase MgdABCD)^47,48^. Other metabolisms encoded by the MAGs include the potential for beta-oxidation of fatty acids, glycogen degradation, fructose and acetate use **(Figure 5, SData15)**. The Pit transporter was identified in all 5 MAGs, as was PhoU and the PhoR-PhoB phosphate regulatory system. The PstABCS system for phosphate transport and the PhnABC system for phosphonate transport were determined in the 4 *Ca*. M. roskildensis MAGs but not *Ca*. M. haderslevensis. In contrast, *Ca*. M. haderslevensis encoded the full pathway for PHA storage (PhaABCZ), whereas the *Ca*. M. roskildensis all missed the *phaB* gene. Nitrogen metabolism also differed between the two species. Nitrogen fixation and respiratory nitrate reduction to nitrite was present in *Ca*. M. roskildensis, but not *Ca*. M. haderslevensis which had periplasmic nitrate reductase (NapAB) and encoded nitrite reduction to nitric oxide and onto nitrous oxide **(Figure 5)**. This suggests different roles or niches for the two species.

### Etymology

Description of *‘Candidatus* Methylophosphatis’ gen. nov. (midas_g_190)

*‘Candidatus* Methylophosphatis’ [Me.thy.lo.phos.pha’tis. N. L. n. methylum (from French *méthyle*, back-formation from French *methylene*, coined from Gr. n. *methu*, wine and Gr. n. *hulê*, wood), the methyl group; N.L. pref, *methylo-*, pertaining to the methyl radical; *phosphatis* from L. nm. phosphas, phosphate: a likely methylotroph which can accumulate polyphosphate].

Description of ‘ *Candidatus* Methylophosphatis roskildensis’ sp. nov. (midas_s_484)

*‘Candidatus* Methlylophosphatis roskildensis’ [ros.kil.den’sis. N. L. fem. adj. roskildensis pertaining to the city of Roskilde, the city in which the Bjergmarken WWTP is situated, and where the sample with the most abundant population was obtained].

Description of *‘Candidatus* Methylophosphatis haderslevensis’ sp. nov. (midas_s_514)

‘Candidatus Methylophosphatis haderslevensis’ (midas_s_514) [ha.der.sle.ven’sis. N. L. fem. adj. haderslevensis pertaining to the city of Haderslev, where the sample was obtained from which the MAG was produced].

## Methods

### Sample processing and DNA extractions

Fresh activated sludge samples were received from 23 Danish WWTP in August and September 2018. Samples were aliquoted into 1.6 mL replicates and frozen at −80°C prior to processing. For extraction, each 1.6 mL tube was decanted into an empty powersoil 2 mL tube. The sample was then spun at 10000 xg for 5 minutes, and most supernatant was discarded. Powersoil beads were transferred back to the tube with the pelleted activated sludge, and the remaining steps followed the manufacturer’s protocol, except vortexing steps were substituted with gentle tube inversion (x10) and the DNA was eluted from the spin column using 60 μl of solution C6. Complementary DNA for differential coverage binning was selected from 2016, and 2017 for the same plants. This DNA was extracted previously using the FastSpin kit as described in^17^. The WWTP Viby was new to the MiDAS program in 2018, so DNA extracted using FastSpin from February and May of 2018 was selected for differential coverage. DNA was temporarily stored in a −20°C freezer until size selection. Size selection of the DNA followed the adapted Oxford Nanopore community SPRI protocol originally adapted from Schalamun and Schwessinger (2017)^49^. DNA concentration and quality was checked using the Agilent TapeStation genomic DNA screen tapes, Qubit 2.0 flurometer (Thermo Fisher Scientific, MA, USA), and Nanodrop ND1000 (Thermo Fisher Scientific, MA, USA). After size selection, DNA was stored in TE buffer (pH 8).

### Library preparation and sequencing

Library preparation for Oxford Nanopore sequencing used the one-pot native barcoding protocol^50^ with the EXP-NBD104 native barcodes, followed by Oxford Nanopore ID Genomic DNA by ligation (SQK-LSK109) (PromethION) according to the manufacturer's instructions. Samples were barcoded and then pooled in triplicate at equimolar concentrations. Pooled samples were run on the Oxford Nanopore PromethION R9 FLO-PRO002 flow cells. If samples did not reach the desired output of at least 20 Gbp of data, they were rerun and the data pooled. Eleven flow cells were used in total, and each flow cell was run within eight weeks of delivery. Flow cells were run for 48 hours.

For Illumina sequencing, genomic DNA was quantified with Qubit 2.0 DNA HS Assay and quality assessed by agarose gel imaging. Library preparation and sequencing of the Illumina data were conducted through Genohub (Austin, TX) and Admera Health (South Plainfield, NJ). Libraries were constructed using a PCR free workflow following the manufacturer’s protocol of NEBNext^®^ Ultra^™^ II DNA Library Prep Kit for Illumina targeting an insert size of 300-400 bp, and run on an Illumina Hiseq X generating 922.5 Gbp of 2 x 150 bp PE data. Library QC was performed with Tapestation D1000 Assay (Agilent Technologies, CA, USA), and quantified using qPCR with QuantStudio^®^ 5 System (Applied Biosystems, USA)**(SData1)**.

### Basecalling and data QC

The 17TB of raw Nanopore fast5 data was basecalled using Guppy v2.2.3 for PromethION and the dna_r9.4.1_450bps_flipflop_prom.cfg model from Oxford Nanopore. 1081 Gbp of data was producedin total from the flow cells after basecalling, and after demultiplexing the yield ranged from 21 – 59 Gbp per sample **(SData1, SData2)**. The data was explored using MinionQC v1.4.0^51^. Samples were demultiplexed, and barcodes and adapter sequences trimmed using qcat v1.0.1 (https://github.com/nanoporetech/qcat). using the following flags -b -k NBD103/NBD104 --trim --detect-middle. Following demultiplexing, the Nanopore reads were further processed using Filtlong v0.2.0 (https://github.com/rrwick/Filtlong) to remove reads <400bp using -min_length 4000 and to remove low quality reads with <80% base call accuracy using -min_mean_q 80. Porechop v0.2.3 (https://github.com/rrwick/Porechop) was used to check the reads for residual barcodes and adapters using default settings and the flag –min_split_read_size 4000, and this was followed by a final round of Filtlong using the described parameters.

The 0.92 Tbp of Illumina data was quality checked using FastQC v0.11.7^52^ and MultiQC v1.7^53^. Cutadapt v1.16^54^ removed the sequencing adaptors, reads <100 bp, reads with a qscore <20, trim ambiguous bases (N) from the start and end, and remove reads with ambiguous base using the parameters -m 100 -q 20 --max-n=COUNT 0 --trim-n.

### Metagenome assembly and polishing

Long-read assembly and polishing following methodology and used GNU parallel v20190122^55^ extensively. The quality processed Nanopore reads were assembled using CANU v1.8^56^ using the parameters: corMinCoverage=0 corOutCoverage=all corMhapSensitivity=high correctedErrorRate=0.105 genomeSize=5m corMaxEvidenceCoverageLocal=10 corMaxEvidenceCoverageGlobal=10 oeaMemory=32 redMemory=32 batMemory=200. If the assembly failed due to set memory limitations, the genome size was increased to -gs 50000000 and the script restarted from the fail point, as suggested by the developers. Once the assemblies were produced, MUMmer v3.2.3^57^ nucmer was used on the contigs CANU labelled as circular (suggestCircular=yes) to show overlaps, so that the duplicated sequence could be removed using R v3.5.0^58^ and unix cut prior to assembly polishing. Contigs <4000bp were removed using seqtk v1.3-r106 (https://github.com/lh3/seqtk) seq. Nanopore assembly polishing was accomplished using Racon v1.3.3 (https://github.com/lbcb-sci/Racon) with the argument --include-unpolished for initial correction based on the mapping of the quality process nanopore reads using minimap2 v2.16^59^ -x map-ont. This was followed by two rounds of medaka v0.6.5 (https://github.com/nanoporetech/medaka). comprising mini align, medaka consensus using the r941_flip213 model and medaka stitch commands. For medaka consensus, the assemblies were split into first 10 (for medaka round 1) and then 20 (for medaka round 2) different files with the contig headers space separated and provided to the --region flag. This was done at the developer’s suggestion in order to improve processing time. Following the second round of medaka, minimap2 was used to map the quality processed Illumina reads to the medaka polished assembly, and Racon --include-unpolished was used again with these reads for a final round of error correction. Samtools v1.9^60^ was used as a dependency for this pipeline.

### MAG binning and dereplication

Binning followed the pipeline mmlong v0.1.2 hybrid-metaflow after the assembly step (https://github.com/SorenKarst/mmlong). Briefly, Nanopore and Illumina reads were mapped to the polished Nanopore assembly using minimap2 v2.15. Automatic binning was conducted using MetaBAT2 v2.12.1^61^ and MaxBin v2.2.7^62^. Metagenome contigs were translated into proteins using FragGeneScan v1.31^63^, annotated taxonomically using Kaiju v1.6.0^64^ against the proGenomes database (2017-05-16), 16S rRNA genes were identified with barmap v0.9 (https://github.com/tseemann/barmap). and then classified with MOTHUR v2.7.14 classify.seqs against the SILVA v132 seed database^65^. Binning was conducted using two coverage approaches for each of the two binning tools, first using the differential coverage information from only the same plant as the assembly (i.e. the corresponding 3 Illumina metagenomes 2016, 2017 and 2018), or differential coverage information from all of the Illumina metagenomes (69 in total). DASTool v1.1.1^66^ --search engine diamond was used to dereplicate and select for the best representative bin from the four binning iterations for each of the 23 metagenomes.

The dereplicated bins were then checked from completeness and contamination using CheckM --lineage_wf^67^ v1.0.11, resulting in 3733 MQ to HQ MAGs. Circular genomes were identified by linking the contig names back to the suggestCircular=yes CANU designations. Circular contigs >700 kbp were identified as likely circular chromosomes. Ultra-small genomes with a circular chromosome were included in the HQ set despite not full-filling the completeness cut-off of >90%.

Five MAGs failed the contamination threshold and were manually examined using mmgenome2 v2.0.7 (https://github.com/KasperSkytte/mmgenome2). four were circular and one was a MAG of interest (*Nitrospira*). Likely contaminating contigs were removed and quality was re-checked with CheckM. These manually improved MAGs are identified by ‘v2’ in the MAG ID in the supplementary tables. Of 43 MAGs containing a circular contig of >700,000 kb, 29 circular MAGs were manually checked and additional non-circular extraneous contigs were removed. These are identified by ‘cln’ in the MAG ID. Three MAGs that contained circular contigs but also large additional linear contigs encoding single copy marker genes, were potentially multichromosomal and were removed from the CMAG designations.

dRep v2.3.2^68^ -comp 50 -con 10 dereplicated the MAGs at 99% ANI clustering to indicate the number of distinct lineages and overlap of likely strains between WWTPs, and at 95% ANI to indicate the number of distinct species.

### Reassembly of 17 Patescibacteria additional CMAGs

While 13 circular Patescibacteria MAGs were initially recovered in the automated binning run with CANU, there were hundreds of additional MQ Patescibacteria MAGs (334) and many had high relative abundances in the metagenomes (e.g. EsbW_18-Q3-R4-48_MAXAC.283, SData3). MQ Patescibacteria MAGs with >30% of their total base pairs in the longest contig were selected for reassembly with Flye v2.6^69^ and Unicycler v0.4.6^70^. First, the Nanopore and Illumina reads that had mapped to bin contigs in the original CANU assemblies were extracted using samtools. These were used as input to Unicycler (unicycler -1-2-1 --no_correct -- min_kmer_frac 0.3 --kmer_count 5 --no_pilon --keep 3 --mode bold --min_fasta_length 1000) and Flye (--nano-raw --genome-size 1m). The Flye and Unicycler assembly graphs were checked in Bandage v0.8.1^71^ to determine which assemblies were now circular. Overall, 11 additional circular Patescibacteria MAGs were recovered with Flye, and 6 additional were recovered using Unicycler. These MAGs are identified by ‘fly’ or ‘uni’ in the MAG IDs. The circular contigs from the Flye assemblies were further polished with minimap2, Racon with Nanopore reads, medaka, and Racon with Illumina reads following the method used for the CANU assembly above, except medaka_consensus was used instead of the mini align, medaka consensus and medaka stitch substeps. The Flye MAGs were checked for circularity by ensuring reads mapped across the contig start and end based on manual inspection in Tablet^72^.

EsbW_18-Q3-R4-48_MAXAC.283 was further refined based on the mappings to include split tandem repeats flanking the viral sequence, which were present in the mapped reads but had been trimmed by the assembly.

Polymorphic rates for all HQ MAGs were determined using CMSeq as described in Pasolli et al. 2019 (https://bitbucket.org/CibioCM/cmseq/). using the poly.py and polymut.py scripts and --mincov 10 --minqual 30 --dominant_frq_thrsh 0.8 **(SData3)**.

### MAG taxonomy and phylogeny

Genome taxonomy was determined using GTDB-Tk v0.3.2^73^ and the refseq release 89 (2019-06-19) database, and the dependencies pplacer v1.1^74^, FastANI v1.2^75^, Prodigal v2.6.2^76^, FastTree 2 v1.2^77^, HMMER v3.1b2^78^ **(SData3)**. Both MQ (MAGs with >50% completeness, <10% contamination) and HQ MAGs were classified. ARB v6.0.3^79^ and IToL v5.5^80^ were used for visualising and refining the tree created by GTDB-Tk, and for illustrating the MAG phyla in **Figure 2**. Pavian^81^ was used to create the Figure 3b Sankey, and is based on the taxonomy assigned by GTDB-Tk.

### Genome annotations and 16S rRNA gene mapping to MiDAS3

Prokka v1.14 --meta --kingdom Bacteria or Archaea^82^ and Infernal v1.1.2^83^ (arguments: cmscan --cut_ga --rfam --nohmmonly --fint 2) were run on the MAGs >90% complete and <5% contamination, and the reduced genome circular MAGs to identify the 16S, 23S, 5S rRNA and tRNA genes. Only the MAGs with full-length rRNA genes and >18 tRNA genes determined with either Prokka and/or Infernal were perceived as high quality and selected for further analyses, partial matches in Prokka or Infernal were discarded. 16S rRNA gene sequences were extracted using BEDTools v2.27^84^ or Fxtract v2.3 (https://github.com/ctSkennerton/fxtract). MAG 16S sequences were mapped against the SILVA v138 nr99 database, and MiDAS ESV and ASV databases using USEARCH v11^85^, firstly orientating the sequences against SILVA with usearch11 -orient, then using -usearch_global -top_hit_only -strandplus -id 0.99 (ASV) -id 0.7 (ESV and SILVA) -maxaccepts 0 -maxrejects 0 -blast6out. The MAG ASV and ESV designations were matched to those in the MiDAS3 database to determine how many of the top 100 species or ASVs had been recovered in a HQ MAG^17^.

EnrichM v0.5.0 (https://github.com/geronimp/enrichM) annotate using Diamond v0.9.22^86^ blasted the MAG proteins against the v10 database, incorporating a KO annotated uniref100 database. EnrichM classify was used to reveal the complete KO modules present in the MAGs. The modules searched for included custom module files from Woodcroft et al. 2018, and this study **(SData12)**. These data were used for the construction of the metabolism Figure 4. The GTDB taxonomy is used for the description of the functional guilds and taxonomy throughout the study, except where the MiDAS3 taxonomy is explicitly stated in the text.

### Mean coverage, relative abundances and recovery success of MAGs

Depth of coverage (mean) based on the Illumina data and Nanopore data was calculated using CoverM v0.3.2 (https://github.com/wwood/CoverM) and the filtered bam files created during the mmlong process described above where the metagenome Illumina and Nanopore reads were mapped directly to the corresponding metagenome assembly **(SData3)**. The following arguments were used: coverm genome -m mean --min-read-aligned-percent 0 --min-read-percent-identity 0 --min-covered-fraction 0. Relative abundances of the MAG species representatives in the metagenomes were calculated by mapping the Illumina data for each of the 69 metagenomes to a concatenated fasta file of the 581 bins using default CoverM settings excepting the following arguments: coverm genome -m relative abundance --min-read-aligned-percent 0.75 --min-read-percent-identity 0.95 --min-covered-fraction 0. Stringent identity and alignment cut offs were used to minimise spurious mappings falsely inflating abundances.

The proportion of the metagenome community recovered in the assembly or the HQ MAG set was investigated using SingleM v0.12.1 (https://github.com/wwood/singlem). SingleM identifies single copy marker genes of 14 ribosomal proteins in short read data, assemblies and bins, and avoids the complications of MAG recovery estimates based on multicopy 16S rRNA genes (https://github.com/wwood/singlem). SingleM pipe was run on the individual metagenomes, assemblies and HQ MAGs. SingleM summarise was then run on the SingleM pipe MAG OTU files to concatenate them into one large table for singlem appraise. This table of OTUs, representing 14 single copy marker genes, could then be compared to the metagenomes with their corresponding assemblies using singlem appraise. This allowed us to determine the percentage of the metagenome community to be recovered at each step. For genus level recovery estimates, the flags --imperfect --sequence_identity 0.89 were used to cluster the OTUs at 89% ANI, or roughly the genus level^7^.

Unbinned populations were also identified using the SingleM data, specifically the OTUs that were abundant in the metagenomes but not matched to the HQ MAG set. Here, SingleM pipe was run with a stringent evalue of ‘le-20’ on the HQ MAGs to avoid spurious hits to homologous regions, and singlem appraise with the flag --output_unaccounted_for_otu_table was used to produce an output table of the unbinned hits (present in the metagenome but not the HQ MAGs). This table was transformed into biom format using singlem summarise --biom_prefix, and these tables were imported into R v3.5.2 using the ampvis2 v2.5.8. R package (https://github.com/MadsAlbertsen/ampvis2). Relative abundance heatmaps of the unbinned populations **(SFig2, SFig3** and **SFig4)** were produced with these data. These data were also used to examine the envOPS12 populations that were not successfully recovered with singlem query. The unbinned marker gene sequences of envOPS12 were used as input to query the MQ and low quality bins of EsbE and EsbW metagenomes.

### *Phylogenetic analysis, probe design and FISH of Paceibacteria* and *Ca*. Methylophosphatis

Phylogenetic analysis of 16S rRNA gene sequences and FISH probe design were performed using ARB v. 6.0.6^79^. Analysis of 16S rRNA gene sequences from the Paceibacteria MAGs was conducted by aligning sequences with MAFFT v7.402^87^, trimming with TrimAL v1.4.rev15^88^ to remove bases with <90% coverage but conserving 60% of the original alignment, and creating a phylogenetic tree in IQ-TREE v1.5.6^89^ using RAxML GTR algorithm with 100 bootstraps. Two FISH-probes, Pac_113 and Pac_683, were used to investigate the lineages. For *Ca*. Methylophosphatis, a phylogenetic tree was calculated based on the aligned 12 full-length ESVs from the genus midas_g_190 and the 16S retrieved from the MAGs, using the PhyML maximum likelihood method and a 1000-replicate bootstrap analysis. Unlabelled competitor probes were designed for single base mismatched non-target sequences. Both sets of probes were validated in silico with mathFISH^90^ to test the hybridization efficiency of target and nontarget sequences. The number of non-target sequences with 0, 1, and 2 mismatches was assessed using the probe match function in ARB. All probes were purchased from Biomers (Biomers.net, Ulm, Germany) and were labelled with ATTO 532 or ATTO 594 fluorochromes (Paceibacteria), or labelled with indocarbocyanine (Cy3) or indodicarbocyanine (Cy5) fluorochromes (*Ca*. Methylophosphatis).

For *Ca*. Methylophosphatis, optimal hybridization conditions for the novel FISH probes were determined based on the formamide dissociation curves generated after hybridization at different formamide concentrations over a range of 0-70% (v/v) with 5% increments. Relative fluorescence intensities of 50 cells were measured with the ImageJ software (National Institutes of Health, Maryland, USA) and calculated average values were compared for selection of the optimal formamide concentration. As a pure culture was not available, the probes were optimized using activated sludge biomass with a high abundance of the target organism predicted by amplicon sequencing. Details about the optimal formamide concentration used for each probe are given in **STable2**.

Fresh biomass samples from full-scale activated sludge WWTP were fixed with 4% paraformaldehyde (final concentration) for 3 hours at 4°C and washed 3 times with 1 mL of sterile filtered tap water, and stored in the freezer (−20°C) until needed. FISH was performed as described by^91^ and^92^ with 3 h hybridization to obtain sufficient fluorescent signal. The EUBmix probe set was used to cover all bacteria^93,94^, and the nonsense NON-EUB probe was applied as negative control for sequence independent probe binding^95^.

Microscopic analysis was performed with either an Axioskop epifluorescence microscope (Carl Zeiss, Germany), equipped with a Leica DFC7000 T CCD camera, or a white light laser confocal microscope (Leica TCS SP8 X) (Leica Microsystems, Wetzlar, Germany). Quantitative FISH (qFISH) biovolume fractions of individual taxa (*Ca*. Methylophosphatis) were calculated as a percentage area of the total biovolume, hybridizing the EUBmix probes, which also hybridises with the specific probe. qFISH analyses were based on 30 fields of view taken at 63× magnification using the Daime image analysis software^96^.

### *Raman microspectroscopy of Ca*. Methylophosphatis (*midas_g_190*)

Raman microspectroscopy was applied in combination with FISH, as previously described^97^, to detect the presence of intracellular storage polymers. Briefly, FISH was conducted on optically polished CaF_2_ Raman windows (Crystran, UK), which give a single-sharp Raman marker at 321 cm^-1^ that serves as an internal reference point in every spectrum. The genus-specific probe for midas_g_190 **(STable2)** was used to locate the target cells for Raman analysis. After bleaching the Cy3 fluorophore with the Raman laser, spectra from single cells were obtained using a Horiba LabRam HR 800 Evolution (Jobin Yvon - France) equipped with a Torus MPC 3000 (UK) 532 nm 341 mW solid-state semiconductor laser. The Raman spectrometer was calibrated prior to obtaining all measurements to the first-order Raman signal of Silicon, occurring at 520.7 cm^-1^. The incident laser power density on the sample was attenuated down to 2.1 mW/μm^-2^ using a set of neutral density (ND) filters. The Raman system is equipped with an in-built Olympus (model BX-41) fluorescence microscope. A 50X, 0.75 numerical aperture dry objective (Olympus M Plan Achromat-Japan), with a working distance of 0.38 mm, was used throughout the work. A diffraction grating of 600 mm/groove was used, and the Raman spectra collected spanned the wavenumber region of 200 cm^-1^ to 1800 cm^-1^. The slit width of the Raman spectrometer and the confocal pinhole diameter were set to 100 μm and 72 μm, respectively. Raman spectrometer operation and subsequent processing of spectra were conducted using LabSpec v6.4 software (Horiba Scientific, France). All spectra were baseline corrected using a 6^th^ order polynomial fit.

### Data availability

Data used in this study, Illumina and Oxford Nanopore metagenomes and HQ MAGs, are deposited in NCBI under the bioproject accession number PRJNAXXXXXX. Both MQ and HQ MAGs have been submitted to Figshare, to enable bulk download under DOI: XXXXXX. Raw FISH images (tif format) are also available in Figshare. Data yield and MAG statistics are presented in the Supplementary Data Files.

## Acknowledgements

We thank the Danish WWTPs for past and present support and participation in the MiDAS project, providing samples for this study. This project was funded by research grants from VILLUM FONDEN (16578, 15510) and the Poul Due Jensen Foundation (Microflora Danica).

## Contributions

P.H.N., M.A., S.M.K., M.S.D., C.M.S., and R.H.K. designed the study. C.M.S., P.H.N. and M.A. wrote the manuscript, and all authors reviewed and approved the final manuscript. C.M.S. and M.H.A. extracted DNA from the samples, carried out the library preparation and sequenced the DNA. S.M.K. and R.H.K. wrote scripts for bioinformatic analysis. C.M.S., R.H.K., and T.Y.M. performed the bioinformatics analyses.

M.S.D. helped with MiDAS data, J.M.K., designed probes, performed FISH and 16S rRNA gene phylogenetic analyses on the Paceibacteria. F.P. designed probes for *Ca*. Methylophosphatis and performed the Raman-FISH analyses and 16S rRNA gene phylogenetic analyses, while Z.W. and F.P. performed probe optimisation and qFISH.

## Ethics declaration

Competing interests

R.H.K., M.A., S.M.K. and P.H.N. own DNASense ApS. The remaining authors declare no competing interests.

## References

1. Tyson, G. W. et al. Community structure and metabolism through reconstruction of microbial genomes from the environment. Nature 428, 37–43 (2004).

2. Venter, J. C. et al. Environmental genome shotgun sequencing of the Sargasso Sea. Science 304, 66–74 (2004).

3. Pasolli, E. et al. Extensive unexplored human microbiome diversity revealed by over 150,000 genomes from metagenomes spanning age, geography, and lifestyle. Cell 176, 649–662.e20 (2019).

4. Shaiber, A. & Eren, A. M. Composite metagenome-assembled genomes reduce the quality of public genome repositories. mBio vol. 10 (2019).

5. Bowers, R. M. et al. Minimum information about a single amplified genome (MISAG) and a metagenome-assembled genome (MIMAG) of bacteria and archaea. Nat. Biotechnol. 35, 725–731 (2017).

6. Stewart, R. D. et al. Compendium of 4,941 rumen metagenome-assembled genomes for rumen microbiome biology and enzyme discovery. Nat. Biotechnol. 37, 953–961 (2019).

7. Woodcroft, B. J. et al. Genome-centric view of carbon processing in thawing permafrost. Nature 560, 49–54 (2018).

8. Ye, L., Mei, R., Liu, W.-T., Ren, H. & Zhang, X.-X. Machine learning-aided analyses of thousands of draft genomes reveal specific features of activated sludge processes. Microbiome 8, 16 (2020).

9. Nielsen, P. H. Microbial biotechnology and circular economy in wastewater treatment. Microb. Biotechnol. 10, 1102–1105 (2017).

10. van Loosdrecht, M. C. M. & Brdjanovic, D. Anticipating the next century of wastewater treatment. Science 344, 1452–1453 (2014).

11. Lawson, C. E. et al. Common principles and best practices for engineering microbiomes. Nat. Rev. Microbiol. 17, 725–741 (2019).

12. Pérez, M. V, Guerrero, L. D., Orellana, E., Figuerola, E. L. & Erijman, L. Time series genome-centric analysis unveils bacterial response to operational disturbance in activated sludge. mSystems 4, (2019).

13. Arumugam, K. et al. Analysis procedures for assessing recovery of high quality, complete, closed genomes from Nanopore long read metagenome sequencing. bioRxiv 2020.03.12.974238 (2020) doi: 10.1101/2020.03.12.974238.

14. Andersen, M. H., McIlroy, S. J., Nierychlo, M., Nielsen, P. H. & Albertsen, M. Genomic insights into Candidatus Amarolinea aalborgensis gen. nov., sp. nov., associated with settleability problems in wastewater treatment plants. Syst. Appl. Microbiol. 42, 77–84 (2019).

15. Gao, H. et al. Genome-centric metagenomics resolves microbial diversity and prevalent truncated denitrification pathways in a denitrifying PAO-enriched bioprocess. Water Res. 155, 275–287 (2019).

16. McIlroy, S. J. et al. MiDAS 2.0: an ecosystem-specific taxonomy and online database for the organisms of wastewater treatment systems expanded for anaerobic digester groups. Database 2017, (2017).

17. Nierychlo, M., Andersen, K. S., Xu, Y., Green, N. & Albertsen, M. Species-level microbiome composition of activated sludge-introducing the MiDAS 3 ecosystem-specific reference database and taxonomy. bioRxiv (2019).

18. Parks, D. H. et al. Recovery of nearly 8,000 metagenome-assembled genomes substantially expands the tree of life. Nat Microbiol 2, 1533–1542 (2017).

19. Luef, B. et al. Diverse uncultivated ultra-small bacterial cells in groundwater. Nat. Commun. 6, 6372 (2015).

20. Parks, D. H. et al. A standardized bacterial taxonomy based on genome phylogeny substantially revises the tree of life. Nat. Biotechnol. 36, 996–1004 (2018).

21. Chen, L. X., Anantharaman, K., Shaiber, A. & Eren, A. M. Accurate and complete genomes from metagenomes. bioRxiv (2019).

22. Lui, L. M., Nielsen, T. N. & Arkin, A. P. A method for achieving complete microbial genomes and better quality bins from metagenomics data. bioRxiv (2020).

23. Sevim, V. et al. Shotgun metagenome data of a defined mock community using Oxford Nanopore, PacBio and Illumina technologies. Sci Data 6, 285 (2019).

24. Schmidt, I. et al. New concepts of microbial treatment processes for the nitrogen removal in wastewater. FEMS Microbiol. Rev. 27, 481–492 (2003).

25. McIlroy, S. J. et al. Identification of active denitrifiers in full-scale nutrient removal wastewater treatment systems. Environ. Microbiol. 18, 50–64 (2016).

26. Ushiki, N. et al. Nitrite oxidation kinetics of two Nitrospira strains: The quest for competition and ecological niche differentiation. J. Biosci. Bioeng. 123, 581–589 (2017).

27. Boddicker, A. M. & Mosier, A. C. Genomic profiling of four cultivated Candidates Nitrotoga spp. predicts broad metabolic potential and environmental distribution. ISME J. 12, 2864–2882 (2018).

28. Morgan-Sagastume, F., Nielsen, J. L. & Nielsen, P. H. Substrate-dependent denitrification of abundant probe-defined denitrifying bacteria in activated sludge. FEMS Microbiol. Ecol. 66, 447–461 (2008).

29. Law, Y., Ye, L., Pan, Y. & Yuan, Z. Nitrous oxide emissions from wastewater treatment processes. Philos. Trans. R. Soc. Lond. B Biol. Sci. 367, 1265–1277 (2012).

30. Chutivisut, P., Isobe, K., Powtongsook, S., Pungrasmi, W. & Kurisu, F. Distinct microbial community performing dissimilatory nitrate reduction to ammonium (DNRA) in a high C/NO3--reactor. Microbes Environ. ME17193 (2018).

31. van den Berg, E. M., Elisário, M. P., Gijs Kuenen, J., Kleerebezem, R. & van Loosdrecht, M. C. M. Fermentative Bacteria Influence the Competition between Denitrifiers and DNRA Bacteria. Frontiers in Microbiology vol. 8 (2017).

32. Onley, J. R., Ahsan, S., Sanford, R. A. & Löffler, F. E. Denitrification by Anaeromyxobacter dehalogenans, a Common Soil Bacterium Lacking the Nitrite Reductase Genes nirS and nirK. Appl. Environ. Microbiol. 84, (2018).

33. McIlroy, S. J. et al. ‘Candidates Competibacter’-lineage genomes retrieved from metagenomes reveal functional metabolic diversity. The ISME Journal vol. 8 613–624 (2014).

34. Nobu, M. K., Tamaki, H., Kubota, K. & Liu, W.-T. Metagenomic characterization of ‘Candidates Defluviicoccus tetraformis strain TFO71’, a tetrad-forming organism, predominant in an anaerobic-aerobic membrane bioreactor with deteriorated biological phosphorus removal. Environmental Microbiology vol. 16 2739–2751 (2014).

35. Oyserman, B. O., Noguera, D. R., del Rio, T. G., Tringe, S. G. & McMahon, K. D. Metatranscriptomic insights on gene expression and regulatory controls in Candidatus Accumulibacter phosphatis. ISME J. 10, 810–822 (2016).

36. Nielsen, P. H., McIlroy, S. J., Albertsen, M. & Nierychlo, M. Re-evaluating the microbiology of the enhanced biological phosphorus removal process. Curr. Opin. Biotechnol. 57, 111–118 (2019).

37. McIlroy, S. J. et al. Genomic and in situ analyses reveal the Micropruina spp. as abundant fermentative glycogen accumulating organisms in enhanced biological phosphorus removal systems. Frontiers in Microbiology 9, (2018).

38. Oshiki, M., Onuki, M., Satoh, H. & Mino, T. PHA-accumulating microorganisms in full-scale wastewater treatment plants. Water Sci. Technol. 58, 13–20(2008).

39. Oshiki, M., Onuki, M., Satoh, H. & Mino, T. Microbial community composition of polyhydroxyalkanoate-accumulating organisms in full-scale wastewater treatment plants operated in fully aerobic mode. Microbes Environ. 28, 96–104 (2013).

40. Nierychlo, M. et al. Candidatus Amarolinea and Candidatus Microthrix are mainly responsible for filamentous bulking in municipal Danish wastewater treatment plants, Frontiers in Microbiology. Front. Microbiol. (Submitted).

41. Albertsen, M. et al. Genome sequences of rare, uncultured bacteria obtained by differential coverage binning of multiple metagenomes. Nat. Biotechnol. 31, 533–538 (2013).

42. Kindaichi, T. et al. Phylogenetic diversity and ecophysiology of Candidate phylum Saccharibacteria in activated sludge. FEMS Microbiol. Ecol. 92, fiw078 (2016).

43. Castelle, C. J. & Banfield, J. F. Major new microbial groups expand diversity and alter our understanding of the tree of life. Cell 172, 1181–1197 (2018).

44. Dueholm, M. S., Andersen, K. S., Petriglieri, F. & McIlroy, S. J. Comprehensive ecosystem-specific 16S rRNA gene databases with automated taxonomy assignment (AutoTax) provide species-level resolution in microbial ecology. Biorxiv (2019).

45. Roux, S., Enault, F., Hurwitz, B. L. & Sullivan, M. B. VirSorter: mining viral signal from microbial genomic data. Peer J 3, e985 (2015).

46. Arndt, D., Marcu, A., Liang, Y. & Wishart, D. S. PHAST, PHASTER and PHASTEST: Tools for finding prophage in bacterial genomes. Brief. Bioinform. 20, 1560–1567 (2019).

47. Smalley, N. E. et al. Functional and genomic diversity of methylotrophic Rhodocyclaceae: description of Methyloversatilis discipulorum sp. nov. Int. J. Syst. Evol. Microbiol. 65, 2227–2233 (2015).

48. Latypova, E. et al. Genetics of the glutamate-mediated methylamine utilization pathway in the facultative methylotrophic beta-proteobacterium Methyloversatilis universalis FAM5. Mol. Microbiol. 75, 426–439 (2010).

49. Schalamun, M. & Schwessinger, B. DNA size selection (> 1kb) and clean up using an optimized SPRI beads mixture, protocols, io. (2017).

50. Quick, J. One-pot native barcoding of amplicons v1 (protocols.io.sg2ebye). protocols.io (2019) doi: 10.17504/protocols.io.sg2ebye.

51. Lanfear, R., Schalamun, M., Kainer, D., Wang, W. & Schwessinger, B. MinIONQC: fast and simple quality control for MinION sequencing data. Bioinformatics 35, 523–525 (2019).

52. Andrews, S. & Others. FastQC: a quality control tool for high throughput sequence data. (2010).

53. Ewels, P., Magnusson, M., Lundin, S. & Käller, M. MultiQC: summarize analysis results for multiple tools and samples in a single report. Bioinformatics 32, 3047–3048 (2016).

54. Martin, M. Cutadapt removes adapter sequences from high-throughput sequencing reads. EMBnet, joumal 17, 10–12 (2011).

55. Tange, O. GNU Parallel 2018. (Lulu.com, 2018).

56. Koren, S. et al. Canu: scalable and accurate long-read assembly via adaptive k-mer weighting and repeat separation. Genome Res. 27, 722–736 (2017).

57. Kurtz, S. et al. Versatile and open software for comparing large genomes. Genome Biol. 5, R12 (2004).

58. R Core Team. R: A language and environment for statistical computing. R Foundation for Statistical Computing. Austria: Vienna. https://www.R-project.org/ (2018).

59. Li, H. Minimap2: pairwise alignment for nucleotide sequences. Bioinformatics 34, 3094–3100 (2018).

60. Li, H. et al. The Sequence Alignment/Map format and SAMtools. Bioinformatics 25, 2078–2079 (2009).

61. Kang, D. D. et al. MetaBAT 2: an adaptive binning algorithm for robust and efficient genome reconstruction from metagenome assemblies. PeerJl, e7359 (2019).

62. Wu, Y.-W., Simmons, B. A. & Singer, S. W. MaxBin 2.0: an automated binning algorithm to recover genomes from multiple metagenomic datasets. Bioinformatics 32, 605–607 (2016).

63. Rho, M., Tang, H. & Ye, Y. FragGeneScan: predicting genes in short and error-prone reads. Nucleic Acids Res. 38, e191 (2010).

64. Menzel, P., Ng, K. L. & Krogh, A. Fast and sensitive taxonomic classification for metagenomics with Kaiju. Nat. Commun. 7, 11257 (2016).

65. Schloss, P. D. et al. Introducing mothur: open-source, platform-independent, community-supported software for describing and comparing microbial communities. Appl. Environ. Microbiol. 75, 7537–7541 (2009).

66. Sieber, C. M. K. et al. Recovery of genomes from metagenomes via a dereplication, aggregation and scoring strategy. Nat Microbiol 3, 836–843 (2018).

67. Parks, D. H., Imelfort, M., Skennerton, C. T., Hugenholtz, P. & Tyson, G. W. CheckM: assessing the quality of microbial genomes recovered from isolates, single cells, and metagenomes. Genome Res. 25, 1043–1055 (2015).

68. Olm, M. R., Brown, C. T., Brooks, B. & Banfield, J. F. dRep: a tool for fast and accurate genomic comparisons that enables improved genome recovery from metagenomes through de-replication. ISME J. 11, 2864–2868 (2017).

69. Kolmogorov, M., Yuan, J., Lin, Y. & Pevzner, P. A. Assembly of long, error-prone reads using repeat graphs. Nat. Biotechnol. 37, 540–546 (2019).

70. Wick, R. R., Judd, L. M., Gorrie, C. L. & Holt, K. E. Unicycler: Resolving bacterial genome assemblies from short and long sequencing reads. PLoS Comput. Biol. 13, e1005595 (2017).

71. Wick, R. R., Schultz, M. B., Zobel, J. & Holt, K. E. Bandage: interactive visualization of de novo genome assemblies. Bioinformatics 31, 3350–3352 (2015).

72. Milne, I. et al. Using Tablet for visual exploration of second-generation sequencing data. Brief. Bioinform. 14, 193–202 (2013).

73. Chaumeil, P.-A., Mussig, A. J., Hugenholtz, P. & Parks, D. H. GTDB-Tk: a toolkit to classify genomes with the Genome Taxonomy Database. Bioinformatics (2019) doi:l0.1093/bioinformatics/btz848.

74. Matsen, F. A., Kodner, R. B. & Armbrust, E. V. pplacer: linear time maximum-likelihood and Bayesian phylogenetic placement of sequences onto a fixed reference tree. BMC Bioinformatics 11, 538 (2010).

75. Jain, C., Rodriguez-R, L. M., Phillippy, A. M., Konstantinidis, K. T. & Aluru, S. High-throughput ANI analysis of 9OK prokaryotic genomes reveals clear species boundaries. bioRxiv. (2017).

76. Hyatt, D. et al. Prodigal: prokaryotic gene recognition and translation initiation site identification. BMC Bioinformatics 11, 119 (2010).

77. Price, M. N., Dehal, P. S. & Arkin, A. P. FastTree 2--approximately maximum-likelihood trees for large alignments. PLoS One 5, (2010).

78. Eddy, S. R. Accelerated Profile HMM Searches. PLoS Comput. Biol. 7, e1002195 (2011).

79. Ludwig, W. et al. ARB: a software environment for sequence data. Nucleic Acids Res. 32, 1363–1371 (2004).

80. Letunic, I. & Bork, P. Interactive Tree Of Life (iTOL) v4: recent updates and new developments. Nucleic Acids Res. 47, W256–W259 (2019).

81. Breitwieser, F. P. & Salzberg, S. L. Pavian: interactive analysis of metagenomics data for microbiome studies and pathogen identification. Bioinformatics 36, 1303–1304 (2020).

82. Seemann, T. Prokka: rapid prokaryotic genome annotation. Bioinformatics 30, 2068–2069 (2014).

83. Nawrocki, E. P., Kolbe, D. L. & Eddy, S. R. Infernal 1.0: inference of RNA alignments. Bioinformatics 25, 1335–1337 (2009).

84. Quinlan, A. R. & Hall, I. M. BEDTools: a flexible suite of utilities for comparing genomic features. Bioinformatics 26, 841–842 (2010).

85. Edgar, R. C. Search and clustering orders of magnitude faster than BLAST. Bioinformatics 26, 2460–2461 (2010).

86. Buchfink, B., Xie, C. & Huson, D. H. Fast and sensitive protein alignment using DIAMOND. Nat. Methods 12, 59–60 (2015).

87. Katoh, K. & Standley, D. M. MAFFT multiple sequence alignment software version 7: improvements in performance and usability. Mol. Biol. Evol. 30, 772–780 (2013).

88. Capella-Gutiérrez, S., Silla-Martínez, J. M. & Gabaldon, T. trimAl: a tool for automated alignment trimming in large-scale phylogenetic analyses. Bioinformatics 25, 1972–1973 (2009).

89. Nguyen, L.-T., Schmidt, H. A., von Haeseler, A. & Minh, B. Q. IQ-TREE: a fast and effective stochastic algorithm for estimating maximum-likelihood phylogenies. Mol. Biol. Evol. 32, 268–274 (2015).

90. Yilmaz, L. S., Pamerkar, S. & Noguera, D. R. mathFISH, a web tool that uses thermodynamics-based mathematical models for in silico evaluation of oligonucleotide probes for fluorescence in situ hybridization. Appl. Environ. Microbiol. 77, 1118–1122 (2011).

91. Wagner, M., Hom, M. & Daims, H. Fluorescence in situ hybridisation for the identification and characterisation of prokaryotes. Curr. Opin. Microbiol. 6, 302–309 (2003).

92. Daims, H., Stoecker, K. & Wagner, M. Fluorescence in situ hybridization for the detection of prokaryotes, in Molecular microbial ecology 208–228 (Taylor & Francis, 2004).

93. Amann, R. I. et al. Combination of 16S rRNA-targeted oligonucleotide probes with flow cytometry for analyzing mixed microbial populations. Appl. Environ. Microbiol. 56, 1919–1925 (1990).

94. Daims, H., Brühl, A., Amann, R., Schleifer, K. H. & Wagner, M. The domain-specific probe EUB338 is insufficient for the detection of all Bacteria: development and evaluation of a more comprehensive probe set. Syst. Appl. Microbiol. 22, 434–444 (1999).

95. Wallner, G., Amann, R. & Beisker, W. Optimizing fluorescent in situ hybridization with rRNA-targeted oligonucleotide probes for flow cytometric identification of microorganisms. Cytometry vol. 14 136–143 (1993).

96. Daims, H., Lücker, S. & Wagner, M. daime, a novel image analysis program for microbial ecology and biofilm research. Environ. Microbiol. 8, 200–213 (2006).

97. Fernando, E. Y. et al. Resolving the individual contribution of key microbial populations to enhanced biological phosphorus removal with Raman-FISH. ISME J. 13, 1933–1946 (2019).

